# Oncolytic HSV-IL27 expression improves CD8 T cell function and therapeutic activity in syngeneic glioma models

**DOI:** 10.1101/2025.05.12.653429

**Authors:** Alexia Martin, Jack Hedberg, Ilse Aguirre-Hernandez, Uksha Saini, Doyeon Kim, Yeaseul Kim, Ravi Dhital, Kevin A. Cassady

## Abstract

**Background:** Malignant gliomas (MG) are the most common primary brain malignancies and are considered universally fatal. Oncolytic HSVs (oHSV) are promising immunotherapeutics capable of selectively lysing cancer cells, eliciting anti-tumor immunity, and providing local delivery of immune-activating transgenes. IL-27 is a pleiotropic cytokine capable of enhancing tumor-reactive cytotoxic T cell (CTL) function while also possessing neuroprotective properties. We hypothesized that IL-27 expression by oHSV would enhance CTL function and improve anti-glioma therapeutic activity.

**Methods:** We developed an oncolytic herpes simplex virus (oHSV) that expresses IL-27 (C027). The anti-glioma efficacy of C027 was tested in three syngeneic orthotopic glioma models derived from both chemical (CT-2A) and genetic (SB28, KR158) glioma lines. Spectral flow cytometry was used to assess immunophenotypic and functional changes in the tumor infiltrates and systemically. To further investigate the C027-related CTL activity, we employed *in vivo* cell specific depletion and IL-27 blockade alongside in vitro T cell stimulation assays. Local and systemic antitumor memory was evaluated by both orthotopic and flank tumor rechallenge of C027-treated long-term survivors.

**Results:** C027 significantly prolonged survival in syngeneic orthotopic glioma models derived from both chemical (CT-2A) and genetic (KR158, SB28) glioma lines. In the CT-2A model, IL-27-expressing oHSV treatment was associated with increased intratumoral multifunctional effector cytotoxic T lymphocytes (CTL) and functional T cell populations systemically. Mechanistically, both CD8 T cells and IL-27 were required for the C027 survival benefit *in vivo* and IL-27 enhanced CTL function *in vitro*. C027-treated mice that survived their initial tumors had local and systemic anti-glioma memory rejecting tumors on rechallenge.

**Conclusions:** Our findings demonstrate that IL-27 expression by oHSV significantly improves anti-glioma therapeutic efficacy, enhances CTL effector function, and induces durable immune memory. Thus, IL-27-oHSV may provide a promising therapeutic approach for malignant gliomas.

- **What is already known on this topic** – Malignant gliomas are highly aggressive tumors largely resistant to current immunotherapies. Oncolytic herpes simplex viruses (oHSV) are promising immunotherapy agents for malignant gliomas and provide a platform for immunomodulatory gene expression.
- **What this study adds** – In this study, we present a novel IL-27 expressing oHSV (C027) that improves survival in syngeneic glioma-bearing mice through a CD8 T cell and IL-27 dependent mechanism and induces durable immune memory.
- **How this study might affect research, practice or policy** – Our study demonstrates that IL-27-expression by oHSV enhances anti-tumor immunity and glioma efficacy suggesting its potential as a novel therapeutic.

## Background

Malignant gliomas (MG), including glioblastomas (GBM), are the most common and aggressive primary malignancy of the central nervous system (CNS).^1^ Prognosis remains poor with a median overall survival of 15 months for patients receiving standard of care involving surgical resection, radiation therapy, and temozolomide for primary GBM.^1–3^ However, with nearly complete recurrence rates, GBMs are effectively incurable.^2, 3^ Tumor location (immunologically distinct CNS), intratumoral heterogeneity, and highly immunosuppressive tumor microenvironment (TME) contribute to therapeutic resistance and poor clinical outcomes.^4, 5^ With little improvement in survival rates over recent decades, novel therapeutics capable of overcoming these challenges are urgently needed to improve outcomes for patients.^5, 6^

Oncolytic viruses present an emerging form of cancer immunotherapy.^7, 8^ We and others have investigated the antitumor efficacy of attenuated oncolytic type I herpes simplex viruses (oHSV) lacking the neurovirulence gene, γ_1_34.5.^9–14^ These engineered viruses can selectively replicate in and lyse cancer cells; however, they are unable to replicate efficiently in post-mitotic cells.^15^ In addition to the direct viral oncolysis, oHSVs can activate the endogenous immune response converting the immunosuppressive “cold” TME toward a more inflammatory “hot” TME.^11, 16^ Further engineering of oHSV with transgenes encoding immunomodulatory molecules (e.g., cytokines, tumor antigens, antibodies) for expression within the TME aim to enhance the anti-tumor immune response.^11, 17–19^

Our recent gene expression analyses from a phase Ib study in which subjects were oHSV-treated with a fractional dose and then underwent resection and injection of the remaining oHSV dose showed that IL27 gene (IL27a/EBI3) expression directly correlated with survival.^13^ Interleukin 27 (IL-27), a member of the IL-6/IL-12 cytokine family, is primarily released by activated antigen presenting cells.^20–22^ The IL-27 receptor (IL-27Rα/gp130) is found on numerous immune and non-immune cell types with highest expression on activated T and NK cells.^20, 22, 23^ IL-27 (p28/Ebi3) is a pleotropic cytokine eliciting both pro- and anti-inflammatory effects in a cell-type and biological context dependent manner.^21^ T cell immunostimulatory effects of IL-27 include enhancing anti-tumor cytotoxic CD8^+^ T lymphocytes (CTLs), T helper cell (Th) type 1 (Th1) CD4^+^ T cell differentiation, and inhibiting Foxp3^+^ T regulatory (T_reg_) cells.^21, 22, 24, 25^ IL-27 can also limit excessive inflammation through inhibition of Th2/Th17 responses, induction of type I T regulatory (Tr1) cells, and stimulation of IL-10 production making it appealing for use in the CNS potentially improving tolerability.^21, 23, 26^ We hypothesized that IL-27 expression by oHSV would enhance CTL function and improve anti-glioma therapeutic activity.

In this study, we sought to identify if the relationship between IL27 expression in oHSV-treated MGs and improved survival observed in the Phase Ib study is associative or causative. We therefore generated a novel oHSV expressing murine IL-27 (**C027**) using the C134 parent virus, which recently completed Phase I enrollment and is in data analysis phase (NCT03657576) [personal communication]. We demonstrate IL-27 expressing oHSV significantly improves survival in three syngeneic murine malignant glioma models over parent virus and vehicle controls. C027 treatment was associated with an increase in multifunctional CTLs within the TME and improved systemic T cell function. Mechanistically, the C027 anti-glioma activity requires CTLs and IL-27 *in vivo* and we show IL27 related CTL functional changes *in vivo* and *in vitro*. Further, C027- treated responder mice resist local and distant tumor rechallenge suggesting development of a functional memory response.

## Methods

### Cell lines and viruses

Murine malignant glioma cell lines syngeneic in C57BL/6 mice used in this study included CT-2A (kindly provided by Dr. Thomas Seyfriend, Boston College), SB28 (Leibniz Institute DSMZ, AC-880), and KR158B ([KR158] courtesy of Dr. Karlyne M. Reilly at the NCI Rare Tumor Initiative, NIH).^27–30^ Murine lines were grown in Dulbecco’s modified eagle medium (DMEM) 2mM L-glutamine 4.5 mg/ml glucose (Corning) supplemented with 10% fetal bovine serum (FBS).

Vero cells (ATCC, Manassas, Virginia, USA) were grown in DMEM 5% bovine growth serum (BGS) and used for virus generation and titration studies.^31^ Cell lines were used at low-passage and confirmed mycoplasma-free (Universal Mycoplasma Detection Kit, 30-1012K, ATCC). Cells were cultured at 37°C and 5% CO_2_.

Lentivirus production was performed as previously described.^31^ Briefly, firefly luciferase (ffluc)-puromycin was inserted into the lentiviral vector and packaged with envelope (pMD.G) and packaging (pCMV.dR8.91) vectors using TransIT-Lenti transfection reagent (Mirus Bio). Viral supernatant was used to transduce CT-2A and KR158 cells followed by puromycin selection.^31^ Luciferase activity was confirmed using ONE-Glo Luciferase Assay (Promega) according to manufacturer’s instructions.

The viruses HSV-1(F), C134, and C154 have been previously described.^9, 32^ In brief, C134 is a Δγ_1_34.5 virus containing the human cytomegalovirus (HCMV) IRS1 gene under control of the HCMV IE promoter in the UL3/UL4 intergenic region to maintain late viral protein synthesis. C154 is a C134-derived virus with GFP encoded in the γ_1_34.5 locus to express Enhanced Green Fluorescent Protein.^11^ C027 was created by co-transfection and homologous recombination as previously described.^33^ It was isolated from among the progeny after co-transfection of the plasmid encoding the MND-promoter driven mIL-27 bicistronic cassette in the γ_1_34.5 locus and PacI digested C154 viral DNA and sequential EGFP-negative plaque selection in Vero cells (**Figure 1A**).^34^

**Figure 1.**
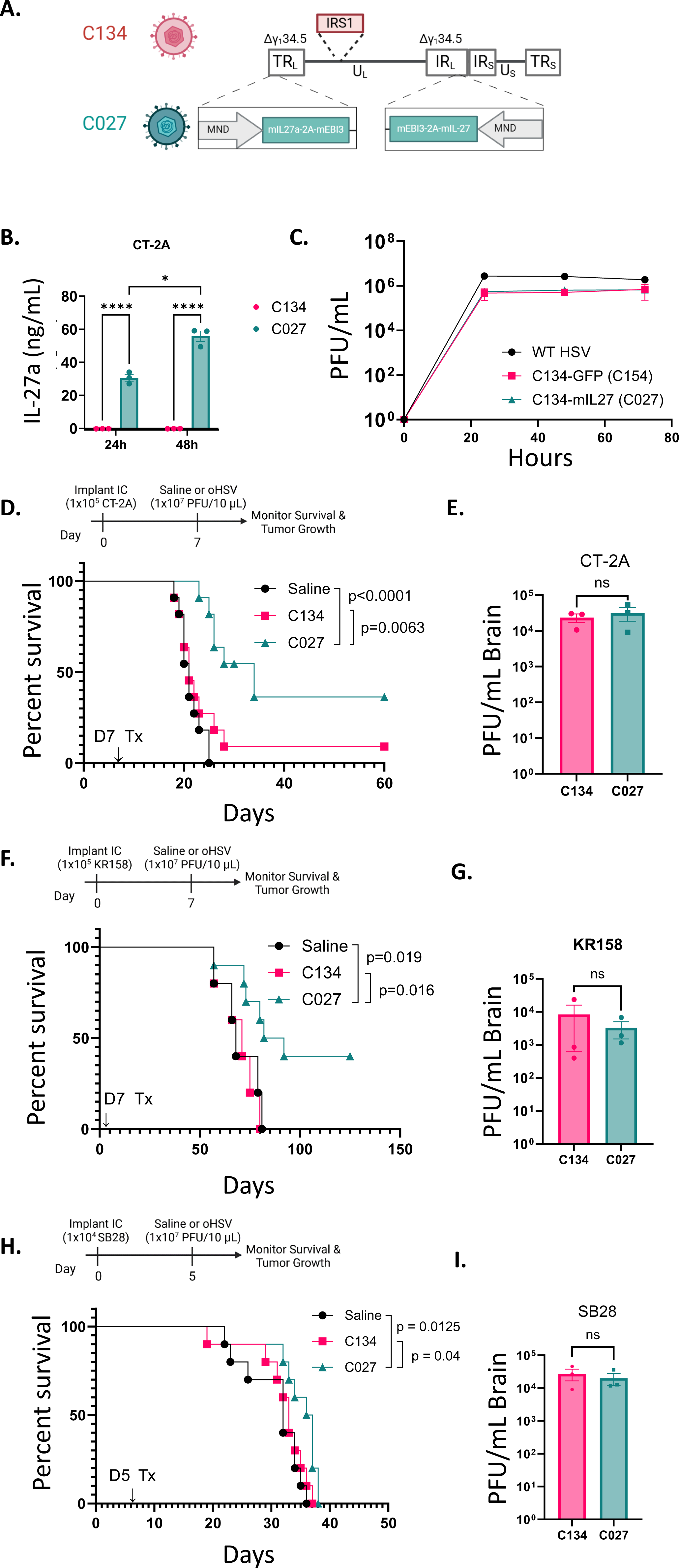
C134-mIL27 (C027) prolongs survival in syngeneic orthotopic murine glioma models. (A) Schematic illustration of C134 and C027. C134 is a second-generation oHSV with deletion of the γ134.5 genes and insertion of the human cytomegalovirus (HCMV) IRS1 gene in the UL3/UL4 intergenic region. C027 was generated from C134 by insertion of bicistronic copies of murine IL-27 (mIL27) in the deleted γ134.5 regions under the MND promoter. (B) mIL27 (p28) secretion from C027 or C134 infected (MOI=1) CT-2A cell supernatants at 24- and 48-hours post-infection measured by ELISA. (C) Viral replication in CT-2A cells infected with C027, C134, or WT HSV (MOI=1) demonstrating viral replication kinetics. Experimental design schematics and representative Kaplan-Meier survival curves in (D) CT-2A (n=11 per cohort), (F) KR158 (n=5- 10 per cohort), and (H) SB28 (n=10 per cohort) immunocompetent syngeneic murine glioma models (Log- Rank test of significance) after treatment with Saline or equivalent PFU (1×10^7^) of C027 or C134 (parental control). *In vivo* viral recovery from murine brain tumors (E) CT-2A, (G) KR158, and (I) SB28 at two days post- treatment with C134 (n=3) or C027 (n=3). For B, C, E, G, and I, data are mean ± SEM with each shape representing one replicate or animal. Statistical analyses were performed using two-way analysis of variance with Holm-Šídák’s correction for multiple comparisons (B), unpaired two-tailed Student’s t-test (E, G, and I), or Log-Rank tests of significance (D, F, and H). IC, intracranial; oHSV, oncolytic herpes simplex virus; PFU, plaque forming unit; MOI, multiplicity of infection; IL, interleukin; Tx, treatment.

### Viral recovery studies

Viral recovery assays were performed as previously described.^31^ Briefly, CT-2A and SB28 cells were plated into 24-well plates and allowed to adhere overnight at 37°C. Cells were infected with equivalent multiplicity of infection (MOI=1) of wild-type, C154 (C134-GFP), or C027 virus for two hours then supernatant removed and replaced with growth medium. Samples were harvested at 24-, 48-, and 72-hours (h) post-infection and viral recovery measured by limiting plaque dilution assay using Vero cells.

### ELISA

Cells were infected with C027 or C154 (MOI=1) and supernatants collected at 24 and 48 hours. Murine IL-27 was quantified using the mouse IL-27 (p28) ELISA kit (Biolegend) according to manufacturer instructions.

### Syngeneic murine malignant glioma models

All animal studies were approved by the Institutional Animal Care and Use Committee (IACUC) at Nationwide Children’s Hospital (AR16-00057, AR21-00145). Female 4-6 weeks old C57BL/6 mice were obtained from Charles River Laboratories and acclimated >3 days prior to study onset. Animals were housed at the Abigail Wexner Research Institute Animal Resources Core under specific pathogen-free conditions.

Tumor cells were harvested during logarithmic growth phase by Accutase (Biolegend) detachment. For intracranial (IC) studies, anesthetized mice were implanted with either 1×10^5^ (CT-2A or KR158) or 1×10^4^ (SB28) in 3 μl of 2.5% methylcellulose using a stereotactic frame (2mm lateral and 1mm anterior to bregma and 2.5mm deep.^10, 11^ Tumor engraftment was confirmed by bioluminescent imaging (BLI) using a Xenogen IVIS Spectrum imaging system then mice were randomized into treatment groups. Five (SB28) or seven (CT-2A or KR158) days (d) post-implantation, anesthetized mice were stereotactically treated with vehicle control or virus (1 x 10^7^ Plaque Forming Units [PFU] / 10 μl) at the same coordinates. Tumor growth was approximated by BLI. Mice were injected subcutaneously with luciferin (D-luciferin sodium salt, 150 mg/kg, GoldBio). Antibodies used for in vivo experiments include: anti-CD8a (2.43, Leinco), anti-IL27 p28 (MM27.7B1, BioXCell), rat IgG2b isotype (LTF-2, BioXCell), mouse IgG2a isotype (N123, Leinco) and were administered by intraperitoneal injection on indicated days. Mice were monitored daily for neurological symptoms and body weight and moribund mice euthanized or at specified time points.

Mice surviving their initial intracranial tumors were rechallenged intracranially in the contralateral hemisphere (1×10^5^ cells) or in the flank with five times the intracranial dose (5×10^5^ cells in 50 μl DMEM / flank) to assess functional antitumor memory responses.^10, 11^ Long-term survivors were defined as surviving >60 days [range 4-10 months] post-initial intracranial implantation with BLI readings at background and rechallenged alongside age-matched naïve controls. Subcutaneous flank tumors were measured by caliper twice weekly until tumor burden exceeded individual (>2000mm^3^) or total (>3000mm^3^) tumor volume was reached.^10, 11^ As per IACUC protocols, mice were euthanized once tumor burden exceeded these criteria. No mice were excluded from analyses.

### Tissue processing and flow cytometry

Tumor-bearing mice were euthanized on specified days post-treatment. Blood was removed by cardiac puncture. Brains were harvested and filtered through a 70 μm cell strainer into a single cell suspension then mononuclear cells isolated by Percoll density gradient (70/30).^11^ Spleens were filtered through a 70 μm cell strainer. Red blood cells were removed from spleens and blood samples using ACK (Ammonium-Chloride-Potassium) lysis buffer.

Tumor infiltrating leukocytes (TIL) and splenocytes were stained and processed as described.^17, 35^ Briefly, cells were incubated with Brefeldin A Solution (BFA, Biolegend, 1X) for 5 hours 37°C. Cells were stimulated with Cell Stimulation Cocktail (eBioscience) for 1h prior to addition of BFA for intracellular cytokine analysis. Dead cells were stained with a fixable viability dye (Zombie NIR Live Dead Stain, Biolegend). After washing, cells were incubated with TruStain FcX™ anti-mouse CD16/32 (Biolegend) and CellBlox blocking buffer (Invitrogen) then stained for surface markers for 45 minutes at room temperature. For intracellular staining, cells were fixed and permeabilized (eBioscience, Foxp3 Transcription Factor Fixation/Permeabilization) following manufacturer’s intracellular staining protocol then incubated with intracellular antibodies overnight at 4°C. Samples were washed and resuspended in FACS buffer for acquisition on a Cytek Aurora spectral flow cytometer using SpectralFlo software version 3.1.0 or Attune NxT Acoustic focusing cytometer using Attune NxT Software v 5.2.0 (Thermo Fischer Scientific). Data were analyzed using FlowJo v 10.8.1 (BD Biosciences) or OMIQ (https://www.omiq.ai/). Detailed information on the complete flow cytometry panel, antibodies, and dilutions used in these studies are provided in **Supplementary Table 1**. Expression of each marker in the spectral flow cytometry panel overlaid on dimenstionality reduced opt-SNE plots are shown in **Supplementary Figure 2** and biaxial gating strategy is detailed in **Supplementary Figure 3**.

Unmixed spectral flow cytometry files (.fcs) were analyzed using the OMIQ platform. Data was arcsinh transformed with cofactor 6000. Manual gating was performed to eliminate debris (FSC v SSC), doublets (FSH v FSC, SSH v SSC), and dead cells (Zombie NIR Live Dead Stain^+^). Total live CD45+ (Zombie NIR Live Dead Stain^-^CD45^+^) and T cells (Zombie NIR Live Dead Stain^-^ CD45^+^ CD19^-^CD3^+^) were manually gated and downsampled 4,000 (TIL live CD45+), 1,500 (TIL T cells), 100,000k (spleen live CD45+), or 10,000 (spleen T cells) events per sample. Data were analyzed using opt-SNE for dimensionality reduction, FlowSOM for clustering, and EdgeR for statistical analyses.^36^ Significant populations identified were also validated by traditional two-dimensional flow cytometry plots.

### In vitro T cell stimulation assays

For T cell stimulation assays, CD3+ T cells were purified from spleens of C57BL/6 mice by negative selection (MojoSort Mouse CD3 T cell Isolation Kit, Biolegend) according to the manufacturer’s instructions. T cells were seeded into 24-well plates in RPMI 10% FBS, 1X antibiotic-antimycotic (Gibco), and IL-2 (10 ng/mL). For stimulated conditions, T cells were activated using soluble anti-CD3 (17A2, Biolegend, 0.5 ug/mL) and anti- CD28 (37.51, Biolegend, 1 ug/mL) with or without murine IL-27 (50 ng/mL). Virally conditioned glioma medium was generated by infecting CT-2A cells (MOI=1) with C027, C134, or vehicle then supernatants were harvested at 24 hours. Splenocytes were isolated from C57BL/6 mice and stimulated as above in the CT-2A conditioned medium for 6 hours. Cells were incubated with BFA for 5 hours prior to collection then processed analyzed by flow cytometry as above (Flow cytometry panel and antibodies used detailed in **Supplementary Table 2**).

### Statistical analysis

Statistical analyses were performed using GraphPad Prism version 10 (GraphPad). Survival data was analyzed by Kaplan-Meier survival curves with log-rank tests of significance. Means between groups were compared using Student’s *t*-tests or one-way or two-way analysis of variance (ANOVA) with Holm-Šídák’s correction for multiple comparisons. Data are represented as mean ± SEM unless otherwise indicated. Statistical significance was defined p values <0.05 (*p<0.05, ** p<0.01, ***p<0.001, ****p<0.0001).

## Results

### Construction and validation of mIL-27 expressing recombinant

Our past phase Ib gene expression studies suggested that IL27 directly correlated with improved subject survival.^13^ To assess whether IL-27 was associative or causative, we constructed an oncolytic HSV expressing murine IL-27, C027 (**Fig 1A**) to test in murine glioma efficacy studies. To evaluate whether C027 produced biologically relevant IL-27 from infected tumor cells, we treated murine glioma cells with C027 (MOI 1) or parent viral control (C134) and measured secreted IL-27 levels from supernatants by ELISA. C027 produced mIL-27 30-47 ng/mL at 24 hours and 47-55 ng/mL at 48 hours (**Fig 1B, Supplemental Fig 1A,B**) and replicated similar to parental virus **(Fig 1C, Supplemental Fig 1C)** in the tested murine glioma cells.

### C027 prolongs survival in syngeneic orthotopic murine glioma models

In order to isolate the immunomodulatory effects of oHSV, we selected three syngeneic murine glioma models. While murine cells are more resistant to sustained oHSV replication and gene expression in vivo, syngeneic models enable examine the endogenous immunotherapeutic effects of oHSV.^10, 17, 37^ To determine if C027 prolongs survival, we initially tested in the carcinogen induced and oHSV resistant CT-2A tumor model.^11^ We implanted CT-2A tumors intracranially (IC) into immunocompetent (C57BL/6) mice and treated intratumorally at the same stereotactic coordinates 7 days later with vehicle or equivalent doses (1×10^7^ PFU) of C134 or C027 (**Fig 1D**). C027 treatment improved survival relative to either vehicle or C134 (median survival; C027 = 34 day vs Saline = 21 days, Log-rank *\*\*\*\*p<0.0001*; C027 = 34 days vs C134= 21 days; Log-rank *\*\*\*p=0.0063*) (**Fig 1D,E, Supplemental Fig 1D**). Consistent with our prior studies, C134 is ineffective in the CT-2A model.^11^ C027 treatment was not associated with overt toxicity with no significant loss of body weight relative to controls (**Supplemental Fig 1E**).

To evaluate whether C027 efficacy was limited to this chemically induced glioma model, we next assessed C027 anti-tumor efficacy in two genetically induced syngeneic murine glioma models (KR158 and SB28). KR158 was derived from a spontaneous tumor in Nf1 and p53 mutant mice whereas SB28 was developed by intraventricular transfection of PDGF-B, NRas, and silencing hairpin-RNA for p53.^28–30, 38^ Similar to the experience with the CT-2A model, C027 again improved survival relative to vehicle (median survival: C027 87d vs. Saline 68d, Log-rank *p=0.019)* and parental virus control, (median survival: C027=87d vs C134=71d; Log rank \**p=0.016*) in KR158 tumors (**Fig 1F,G, Supplemental Fig 1F**). C027 also modestly improved median survival in the SB28 model (Median survival 36.5d vs Saline 31d and C134 32d; Log-rank *p=0.0125*; *p=0.04*), however it did not produce long-term survivors in this model (**Fig 1H,I, Supplemental Fig 1G**). Consistent with the results in the CT-2A model, treatment with parent virus (C134) did not prolong survival relative to vehicle control in both the KR158 and SB28 models.

### IL-27 expression by oHSV increases multifunctional effector CD8 T cells in the tumor microenvironment

After establishing that oHSV-IL27 improved survival in both chemically and genetically derived syngeneic glioma models, we next sought to investigate the mechanism for this anti-glioma therapeutic activity. We first examined oHSV IL-27 immunophenotypic changes in tumor infiltrating leukocytes (TIL) by spectral flow cytometry (day 12 post treatment). As described in the methods section, TILs were isolated from mice bearing intracranial CT-2A tumors 12d post C027 or control (vehicle or C134) treatment and stained using our spectral flow cytometry panel (**Fig 2A**). The results are summarized in **Figure 2** with multi-parameter immune data depicted as immune cell metaclusters (FlowSOM) visualized as reduced dimensionality plots (opt-SNE) (**Fig 2B**). Significant population changes were identified and are represented in volcano plots (**Fig 2C**). Expression of markers on TILs were mapped onto opt-SNE plots (**Supplementary Fig 2**). Gating strategy (**Supplementary Figure 3**), major immune populations (**Supplementary Fig 4,5**) and significant clusters (**Supplementary Fig 6**) were confirmed using traditional two-dimensional dot plot gating.

**Figure 2.**
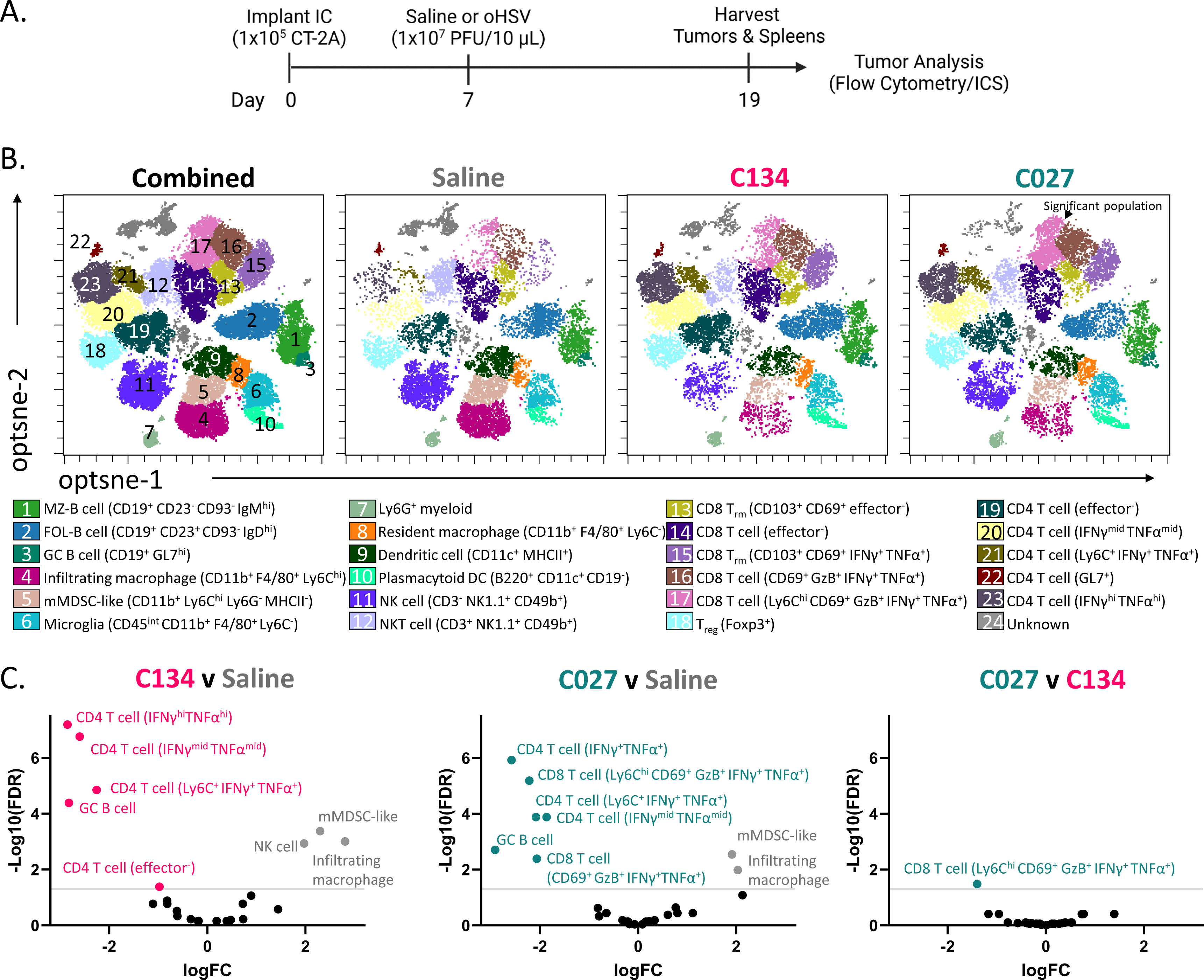
IL-27-oHSV related tumor infiltrating leukocyte immunophenotypic changes. Immunophenotyping of CT-2A murine gliomas treated with vehicle, C134, or C027. tumors were collected and immune cells isolated for spectral flow cytometry analysis 12d post-treatment. (A) Experimental designs schematic. (B) FlowSOM metaclusters overlaid on opt-SNE of all concatenated samples and by treatment cohort using the spectral flow cytometry panel. Each dot represents a single cell. (C) Volcano plots comparing abundance of immune metaclusters between vehicle (gray), C134 (pink), and C027 (teal). Gray line represents a P-value cut-off of 0.05 by EdgeR analyses with statistically significant clusters indicated with cluster labels in volcano plots. N=4 mice per group. IC, intracranial; oHSV, oncolytic herpes simplex virus; PFU, plaque forming unit; ICS, intracellular cytokine staining; MZ-B, marginal zone B cell; FOL-B, follicular B cell; GC B, germinal center B cell; EFF, effector; mMDSC, monocytic myeloid derived suppressor cell; DC, dendritic cell; NK, natural killer; NKT, natural killer T; TRM, tissue-resident memory T cell; Treg, Regulatory T cell; GzB, granzyme B; IFNγ, interferon gamma; TNFα, tumor necrosis factor alpha; FDR, false discovery rate; FC, fold change.

Consistent with our past studies, virus treatment increased the CD45+ populations, enhancing T cell numbers and activity (both activated [CD69^+^ effector^-^], effector cytokine producing [IFNγ^+^ TNFα^+^], and inflammatory [Ly6C^+^] CD4 T cell populations) relative to vehicle treated mice **(Fig 2, Supplemental Fig 4A-B, 6A-C)**. In addition, virotherapy increased germinal center B cells (CD19^+^ GL7^hi^) and activation marker expression (CD69, CD44) compared to saline treated tumors **(Fig 2, Supplemental Fig 6D)**.

In contrast, vehicle treated mice had increased monocytic myeloid-derived suppressor (mMDSC)-like cells (CD11b^+^ Ly6C^hi^ Ly6G^-^ MHCII^-^ F4/80^-^) and infiltrating macrophages (CD11b^+^ F4/80^+^ Ly6C^hi^) (**Fig 2, Supplemental Fig 4C-E, 6E**). Neither viral treatment increased Foxp3^+^ T regulatory cell proportions (**Fig 2B**).When C027 was compared to parent virus treated mice, C027 treatment uniquely increased multifunctional CD8 T cell populations (Ly6C^hi^ CD69^+^ GzB^+^ IFNγ^+^ TNFα^+^). This CD8 population was characterized by co-expression of multiple effector cytokines and was increased in C027-treated tumors relative to both C134- and vehicle-treated mice (**Fig 2B,C**).

Because oHSV, and specifically C027, produced T cell related changes in the tumor microenvironment, we repeated subcluster analysis (opt-SNE, FlowSOM) on the CD3 T cell subpopulation (**Fig 3A**). Consistent with the global CD45+ immune analysis, C027-treated mice demonstrated an in increase in this multifunctional CD8 T cell population relative to parent and vehicle controls (**Fig 3A-C**). In addition to this population, we also identified an increase in non-activated (CD69^-^) and cytokine low (TNFα^lo^ IFNγ^lo^) CD4 T cell populations in the C134-treated mice compared to C027 (**Fig 3A-C**). Relative to vehicle, C027-treatment increased functional tissue-resident memory (T_rm_) CD8 T cells (CD103^+^ CD69^+^ Ly6C^hi^ GzB^+^ IFNγ^+^ TNFα^+^) and functional CD4 T cells (IFNγ^hi^TNFα^hi/mid^) with decreased proportions of non-activated (CD69^-^) and non-functional (effector cytokine^-^) CD8 T_rm_, CD8 T cells, and CD4 T cells (**Fig 3A-C**). In confirmation with the high parameter analyses, standard flow cytometry gating demonstrated increased dual GzB^+^ IFNγ^+^ CD8 and CD4 T cells (**Fig 3D,E**). Overall, C027 treatment increased activated, functional T cell populations and specifically a multifunctional CD8 T cell population.

**Figure 3.**
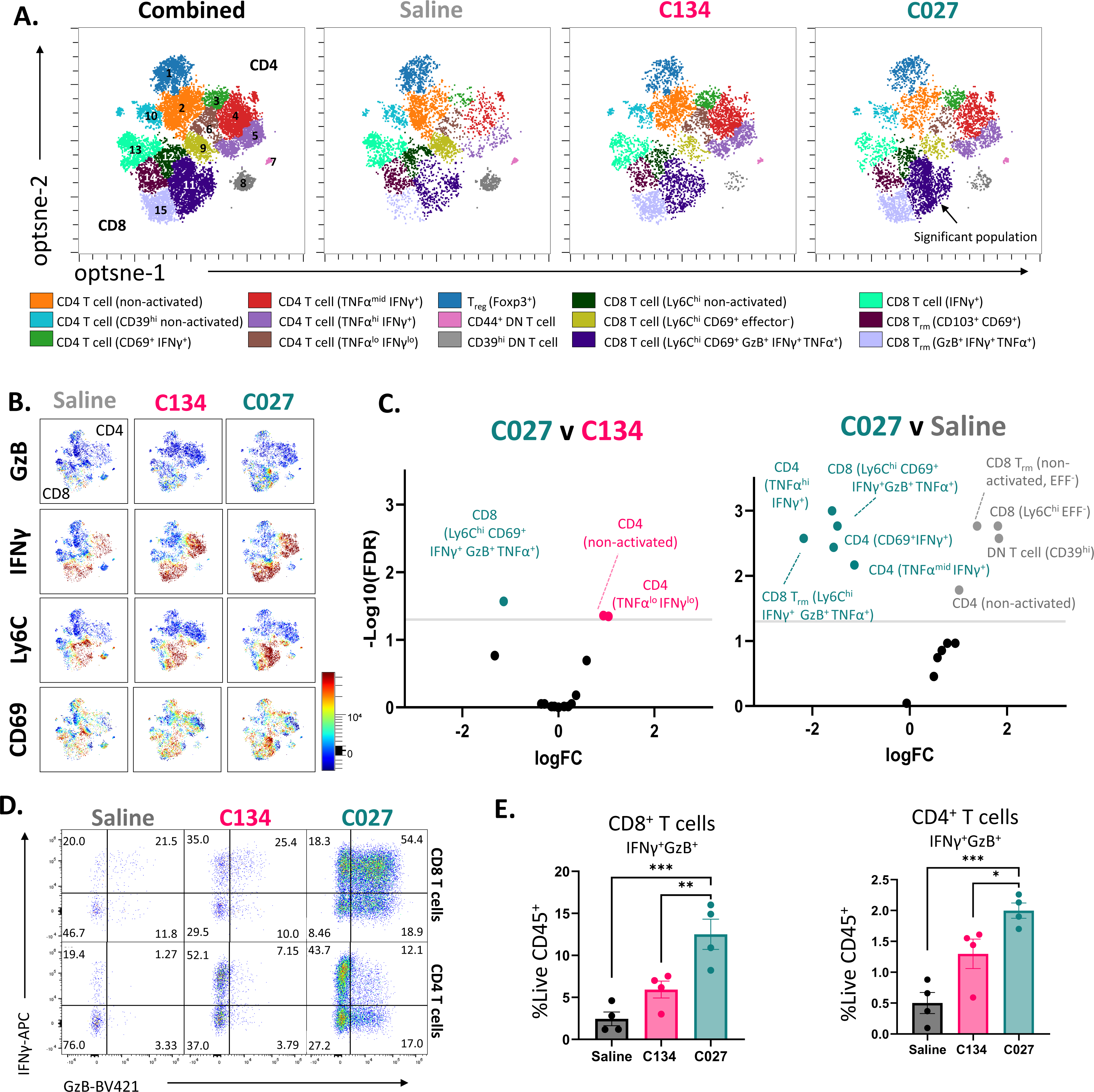
C027 increases multifunctional CD8 effector T cells in the tumor microenvironment. CD3+ T cell sub analyses of CT-2A murine gliomas treated with vehicle, C134, or C027. Twelve days after treatment, tumors were collected and immune cells isolated for spectral flow cytometry analysis (refer to experimental schematic **Fig 2A**). (A) FlowSOM metaclusters of CD3+ T cells overlaid on opt-SNE embedding for all concatenated samples and by treatment cohort. Each dot represents a single cell. (B) Granzyme B (GzB), interferon-gamma (IFNγ), Ly6C, and CD69 expression overlaid on opt-SNE plots of samples concatenated by treatment cohort. Color gradient correlates with marker expression level. (C) Volcano plots comparing proportional CD3+ T cell metaclusters differences between C027 (teal) and vehicle (gray) or C134 (pink). Gray line represents a P-value cut-off of 0.05 by EdgeR analyses with statistically significant clusters indicated with cluster labels in the volcano plots. (D) Representative biaxial flow plots of IFNγ and GzB expression in tumor-infiltrating CD4 and CD8 T cells. (E) Frequencies of dual IFNγ+ GzB+ CD8+ and CD4+ T cells among total CD45+ cells. Statistical analysis was performed by one-way analysis of variance with Holm-Šídák’s multiple comparisons test. N=4 mice per group. Treg, Regulatory T cell; DN, dual CD4 and CD8 negative; TRM, tissue-resident memory T cell; GzB, granzyme B; IFNγ, interferon gamma; EFF, Effector; TNFα, tumor necrosis factor alpha; FDR, false discovery rate; FC, fold change.

### CD8 T cells are required for the anti-glioma efficacy of C027

Next, to determine if the CD8 T cell changes observed in the C027 treated mice were functionally relevant, we repeated orthotopic CT-2A experiments in C57BL/6 mice who underwent selective CD8 depletion or treated with isotype control (**Fig 4A**). The results show that depletion of CD8 T cells eliminated the C027 survival benefit in CT-2A glioma bearing mice (**Fig 4A, Supplementary Fig 1H**). Thus, CD8 T cells are required for the C027 therapeutic activity.

**Fig 4.**
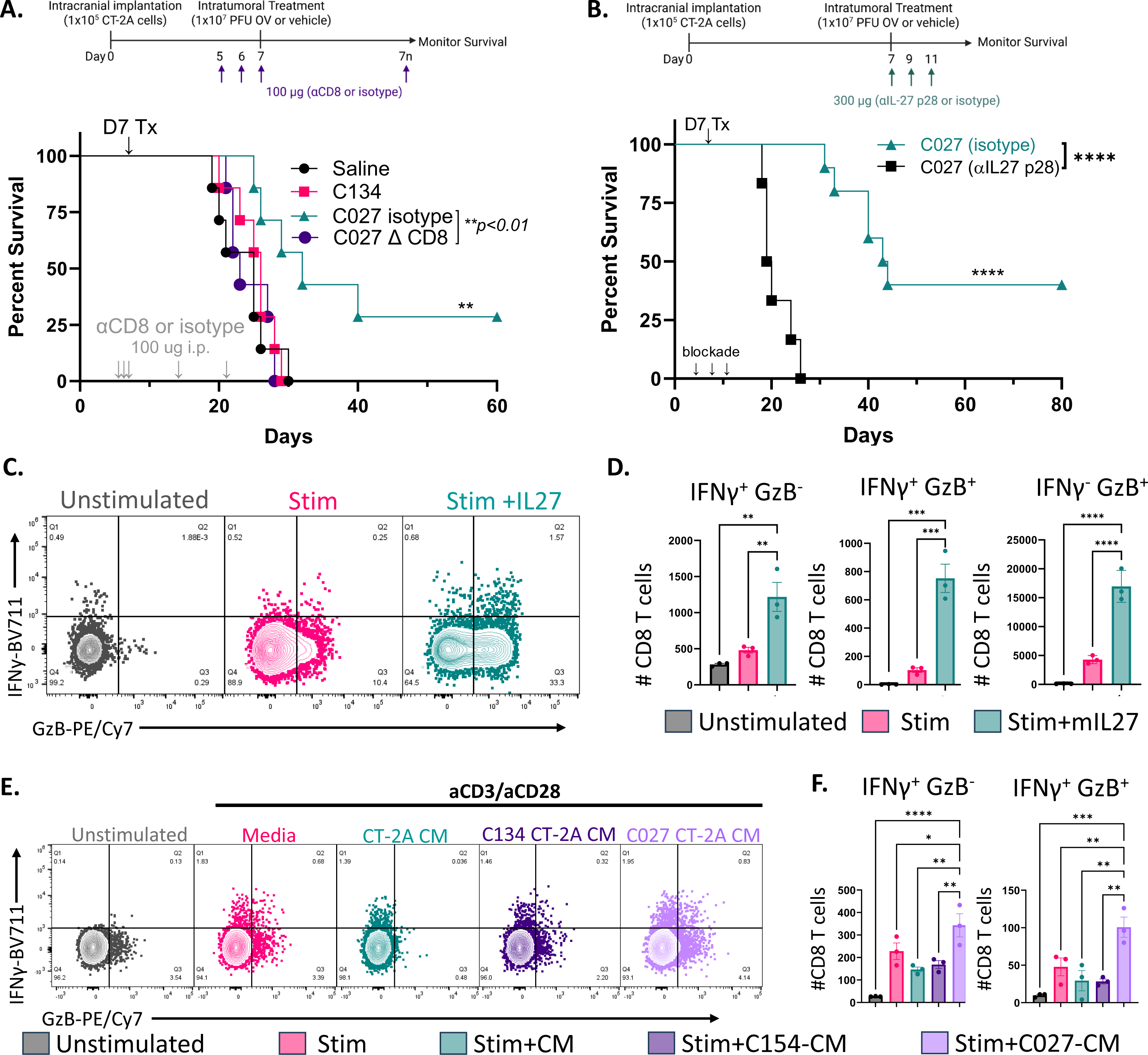
C027 anti-glioma efficacy is dependent on CD8 T cells and IL-27. (A) Experimental design schematic and Kaplan-Meier survival curve of orthotopic CT-2A-bearing mice depleted of CD8 T cells by intraperitoneal injection (days 5, 6, 7, 14, 21) of anti-CD8 or isotype control (n=7 per cohort). (B) Experimental design schematic and Kaplan-Meier survival curve of orthotopic CT-2A-bearing, C027-treated mice receiving intraperitoneal injection (days 7, 9, 11) of IL-27p28 neutralizing antibody or isotype control (n=6-10 per cohort). (C-D) CD3^+^ T cells were stimulated (αCD3/αCD28) +/- IL-27 (50 ng/mL) for 16 hours. (E-F) Splenocytes were stimulated with CT-2A conditioned media (Mock, C154, or C027) for 6 hours. (C, E) Representative flow plots of IFNγ and GzB expression by CD8^+^ T cells. (D, F) abundance of IFNγ^+^ and GzB^+^ CD8^+^ T cells. Data are mean ± SEM with each shape representing one replicate. Statistical analyses were performed by Log-Rank tests of significance (A, B) or one-way analyses of variance with Holm-Šídák’s correction for multiple comparisons (D, F). OV, oncolytic herpes simplex virus; PFU, plaque forming unit; Tx, treatment; i.p., intraperitoneal; stim, stimulated; CM, conditioned media; GzB, granzyme B; IFNγ, interferon gamma.

While C027 significantly extended median survival in CT-2A by 62% relative to vehicle (**Fig 1D**), C134 did not (21d v 21d vehicle) suggesting IL-27 expression is essential for the anti-glioma efficacy. To assess if oHSV-IL27 expression was required for C027 survival benefit, we performed IL-27 blockade studies. In CT-2A-bearing mice who underwent IL-27 neutralization by intraperitoneal injection (anti-IL27 p28 or isotype control), blockade of IL-27 abrogated the C027 survival benefit (Log-rank *\*\*\*\*p<0.0001*) indicating the C027 anti-glioma effects are IL-27 dependent (**Fig 4B, Supplementary Fig 1I**).

Next to further investigate the effects of IL-27 on CD8 T cell activation and effector cytokine production, we stimulated T cells using αCD3/CD28 to mimic activation via CD3/TCR complex recognition of the peptide-MHC complex with or without IL-27. T cells were then stained and analyzed by flow cytometry. Similar to our *in vivo* studies, T cells stimulated in the presence of IL-27 had increased effector cytokine production (IFNγ, GzB) and multifunctional (dual IFNγ^+^GzB^+^) CD8 T cells (**Fig 4C,D**) similar to our *in vivo* observations. To examine the effects of IL-27 in the context of viral production in glioma cells to more closely mimic the *in vivo* system, we generated virally condition glioma medium by infecting glioma cells then collecting the supernatant. Leukocytes were harvest from spleens and activated in the presence of conditioned media and stained for flow cytometry. Similarly, CD8 T cells stimulated in the presence of C027 CT-2A conditioned medium had increased multifunctional (dual IFNγ^+^ GzB^+^) populations (**Fig 4E,F**). These data suggest that IL-27 improves the function of activated CD8 T cells. Taken together, C027 therapeutic activity requires both CD8 T cells and IL-27 wherein IL-27 enhances CD8 T cell function.

### C027-treated long-term survivors resist tumor rechallenge

Both local and systemic immunosuppression occurs in humans and mice with malignant gliomas.^39^ Based on the observed increase in multifunctional T cell populations within the tumor microenvironment, we next examined whether more functional T cell populations were present in the C027-treatment cohort systemically. Splenocytes were isolated from treated (vehicle, C134, or C027) CT-2A bearing mice on day 19 and stained for flow cytometry (**Fig. 5A**, **Supplementary Table 1**). As before, subcluster analysis (opt-SNE, FlowSOM) on the CD3^+^ T cell subpopulation was performed (**Fig 5B**) and significant populations confirmed by traditional 2D gating (**Supplementary Fig 3,7**).

**Figure 5.**
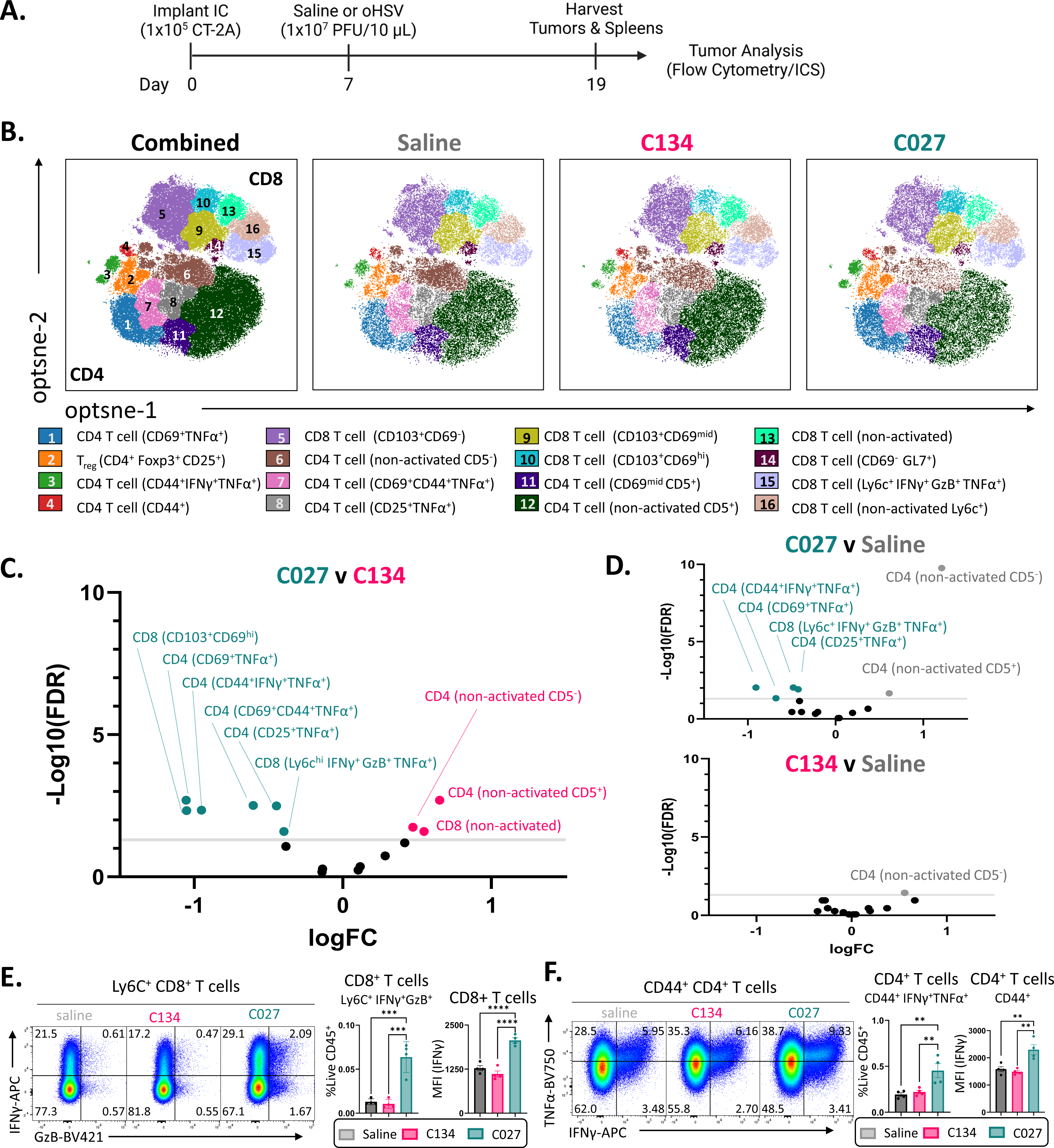
IL-27 expression by oHSV increases functional T cells systemically. Tumor bearing mice were sacrificed 12d post treatment (vehicle, C134, or C027) and their splenocytes analyzed by spectral flow cytometry to assess systemic immunophenotypic changes. (A) Experimental designs schematic. (B) FlowSOM metaclusters overlaid on opt-SNE embedding using the spectral flow cytometry panel. Plots show total concatenated samples and by treatment cohort. Each dot represents a single cell. (C, D) Volcano plots comparing abundance of immune metaclusters between vehicle (gray), C134 (pink), and C027 (teal). Gray line represents a P-value cut-off of 0.05 by EdgeR analyses with statistically significant clusters indicated with cluster labels in volcano plots. (E) Biaxial flow plot of Ly6C+ CD8+ T cells concatenated by treatment cohort. Frequency of IFNγ+ GzB+ Ly6C+ CD8+ T cells and IFNγ+ expression in Ly6C+ CD8+ T cells. (F) Biaxial flow plot of CD44+ CD4+ T cells concatenated by treatment cohort. Frequency of IFNγ+ TNFα+ CD44+ CD4+ T cells and IFNγ+ expression in CD44+ CD4+ T cells. Statistical analyses were performed by one-way analyses of variance with Holm-Šídák’s correction for multiple comparisons. N=4 mice per group. IC, intracranial; oHSV, oncolytic herpes simplex virus; PFU, plaque forming unit; ICS, intracellular cytokine staining; Treg, Regulatory T cell; GzB, granzyme B; IFNγ, interferon gamma; TNFα, tumor necrosis factor alpha; FDR, false discovery rate; FC, fold change; MFI, mean fluorescence intensity.

C027-treated mice had increased activated (CD69^+^, CD25^+^, CD44^+^) and functional (GzB^+^ IFNγ^+^ TNFα^+^) CD4 and CD8 T cell populations with a concomitant decrease in non-activated, non-functional CD4 and CD8 T cell populations relative to parent virus (C134) and vehicle treated mice (**Fig 5B-D**). In validation with the high parameter analyses, standard flow cytometry gating demonstrated increased proportions of multifunctional CD8 (Ly6c^hi^ IFNγ^+^ GzB^+^ TNFα^+^) and CD4 (CD44^+^ IFNγ^+^TNFα^+^) T cells as well as cytokine/activation marker expression in C027-treated mice (**Fig 5E,F and Supplemental Fig 7**). In addition to functional systemic T cell changes, we sought to examine changes in CD8 T cell memory populations. Blood from treated (vehicle, C134, or C027) CT-2A bearing mice on day 19 was collected and stained for standard flow cytometry. C027- treated mice had increased proportions of circulating CD8 T cells, CD8 central memory T cells (CD62L^+^ CD44^+^), GzB^+^ CD8 T cells, and T-bet^+^ CD8 T cells (**Supplementary Figure 8**). Thus, C027 therapy is associated with more functional T cell populations both locally within the tumor and systemically.

We next asked whether C027-treatment elicits a functional memory response capable of protecting against local intracranial tumor rechallenge. C027-treated, long-term survivors ([LTS] >60 days [range 4-10 months] post-initial intracranial implantation, BLI at background levels) were IC rechallenged in the contralateral hemisphere with 1×10^5^ CT-2A cells alongside naïve age-matched controls (**Fig 6A)**. Rechallenged C027-long term survivors rejected their intracranial tumors compared to naïve controls which all succumbed to their tumors (Log-rank *\*p=0.0132*) suggestive of a durable immune memory response (**Fig 6B,C**).

**Figure 6.**
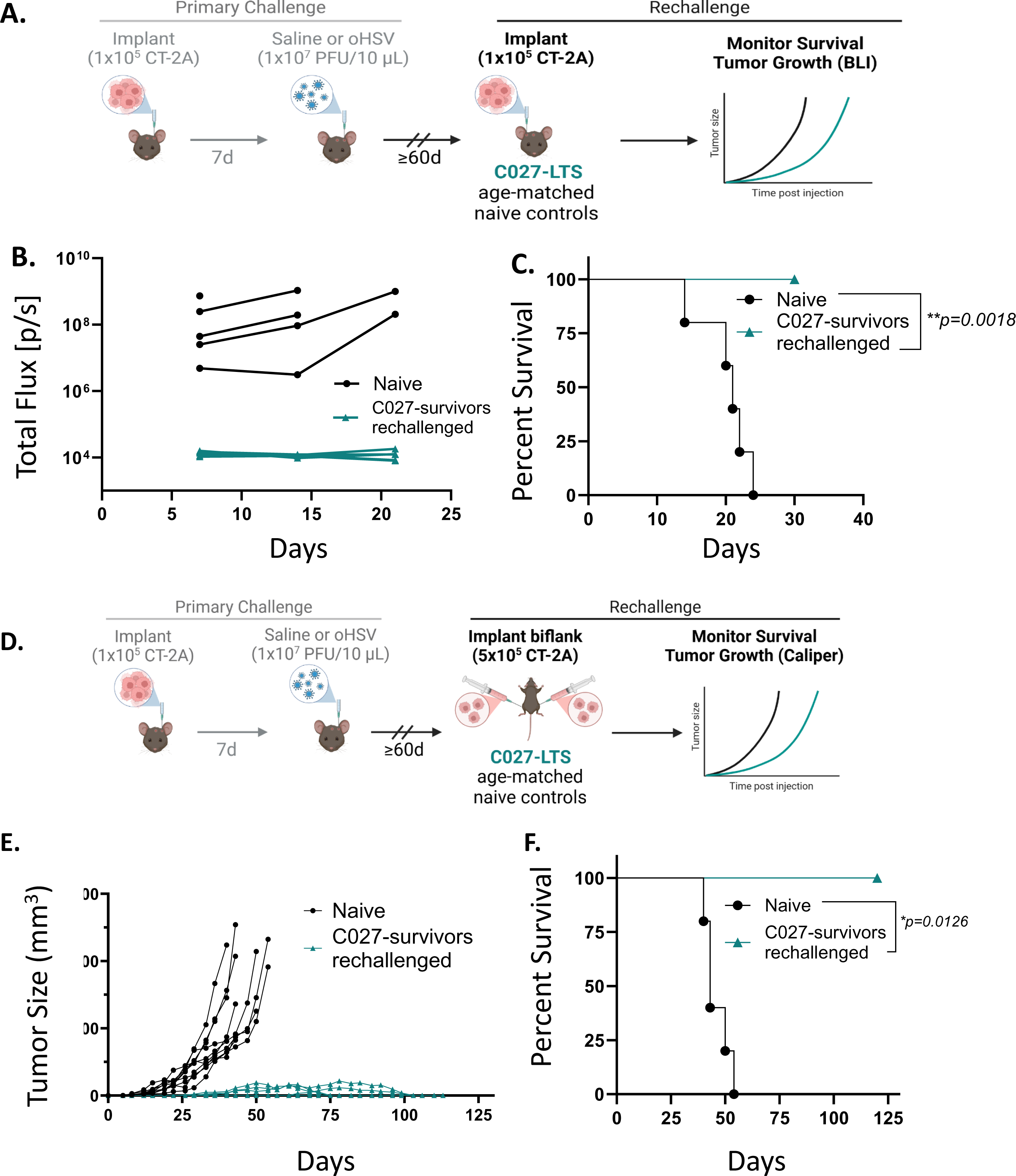
C027-treated responders resist tumor regrowth on rechallenge suggesting development of a durable immune memory response. (A) Schematic of experimental approach. CT-2A-bearing, C027-treated long-term survivors (LTS) were rechallenged intracranially alongside age-matched naïve controls. (B) Tumor growth curves approximated by bioluminescent imaging and (C) Kaplan-Meier survival curves of C027-LTS (teal) and naïve controls (black) (n=5 per cohort). (D) Schematic of experimental approach. CT-2A-bearing, C027-treated long-term survivors (LTS) were rechallenged in the flank alongside age-matched naïve controls. (E) Representative tumor growth curves and (F) Kaplan-Meier survival curves of C027-LTS (teal) and naïve controls (black) (n=3-5 per cohort). Statistical analyses were performed using Log-rank tests of significance. PFU, plaque forming unit; LTS, long-term survivors.

To investigate whether C027 may be producing systemic functional immune memory providing protection from tumor challenge at distant sites, we performed flank rechallenge studies. C027-long-term survivors were rechallenged in the flank with five times the intracranial dose (5×10^5^) of CT-2A glioma cells alongside age-matched naïve controls and tumor growth was measured overtime (**Fig 6D)**. Flank rechallenged C027-long term survivors prevented tumor growth relative to age-matched naïve controls (Log-rank \**p=0.0126*) (**Fig 6E,F**). These results suggest that C027 elicits a functional systemic memory response providing protection from subsequent tumor rechallenge.

## Discussion

Oncolytic HSVs are an encouraging therapeutic approach for treating malignant gliomas through both direct oncolysis and indirect immune activation and modulation of the tumor microenvironment.^7, 8, 13, 16^ More recently, oHSVs are being engineered to enhance these immunostimulatory effects through expression of immune-modulating factors.^11, 17–19^ Further, oHSVs can provide targeted delivery of these molecules for local expression within TME. This localized production provides multiple potential benefits including avoiding possible systemic toxicity, improved transgene expression by virtue of viral expression, and bypassing the blood brain barrier.^15, 16, 18, 40^ Several such viruses have been found safe in early phase I trials for recurrent GBM.^8, 12, 14^ We previously reported on gene expression results from a prior early generation oHSV phase Ib study in which subjects with recurrent GBMs were treated with oHSV followed by resection and found IL-27 gene expression in the oHSV-treated tumors correlated with survival.^13^

In this study, we constructed a novel IL-27 expressing oHSV (C027) aiming to enhance the endogenous T cell response improving the anti-glioma effects. Importantly, C027 demonstrated improved therapeutic activity *in vivo* significantly prolonging survival in three syngeneic murine glioma models relative to parent and vehicle controls. While C027 provided substantial prolongation of median survival in the CT-2A and KR158 models, the C027 effect was marginal in the SB28-gliomas (Median survival 36.5d vs Saline 31d) and failed to generate any long-term responders. This may be due to key differences between the models including lack of MHCI expression in SB28 whereas CT-2A and KR158 have basal MHCI expression.^38, 41^ Further inquiry into the role of tumor-intrinsic MHCI for mediating these effects is needed.

To investigate underlying mechanisms for this effect, we performed immunophenotyping on CT-2A-bearing mice and found C027-treatment was associated with an increase in multifunctional CTLs within the tumors exhibiting a highly activated phenotype and production of numerous effector cytokines. Prior studies have shown multifunctional CTLs have enhanced cytotoxicity and have been associated with effective immune protection and immunotherapeutic response.^42, 43^ One study in patients with metastatic melanoma found expansion of these multifunctional CTLs following dendritic cell vaccination was associated with prolonged survival.^44^

Based on these findings, we hypothesized that the multifunctional CD8 T cells may contribute to the C027 anti-glioma activity and IL-27 may be enhancing their activity. Indeed, the CD8 T cell depletion studies demonstrated that the C027 therapeutic effect required CD8 T cells. Likewise, the C027 survival benefit was completely abrogated in mice treated with IL-27 neutralizing antibody. Similar to the increased multifunctional CTLs observed in C027-treated mice *in vivo*, we show that IL-27 enhances CTL effector cytokine production *in vitro*. Thus, IL-27 expression by oHSV elicits anti-glioma activity in a CD8 T cell and IL-27 dependent manner. Wherein, IL-27 enhances CTL effector function. In agreement with these findings, a recent study published during the preparation of this manuscript found IL-27 expression correlated with CTL gene expression signatures in tumors and immunotherapy response.^45^ Similar to our previous study demonstrating a correlation between IL-27 gene expression in oHSV-treated tumors and survival, the authors found CTL signature was associated with *IL27* expression in multiple tumor types including glioblastoma.^13, 45^ Further, IL-27 overexpression or treatment with engineered IL-27 protein resulted in tumor regression in murine *in vivo* models.^45^ Mechanistically, the authors demonstrate IL-27 acts on CTLs to enhance their cytotoxic function and persistence improving tumor control and activity during chronic antigen stimulation.^45^

Notably, C027-treated mice had increased functional T cells systemically capable of producing multiple effector cytokines upon stimulation. This may be due to a reduction in systemic immune suppression associated with malignant glioma progression or C027-treatment related enhancement of T cell function. While the multifunctional CD8 T cell population was the main immune population identified as unique to C027-treated mice on comparison to C134-treated within the tumor, increases in functional CD4 T cell populations were observed systemically. Further studies are needed to elucidate the role of CD4 T cells in the C027 treatment effect.

In addition, increased circulating CD8 T_cm_ populations were observed in C027-treated mice. Moreover, C027- treated mice cured of their intracranial CT-2A tumors prevented tumor growth on intracranial tumor rechallenge in the contralateral hemisphere. Likewise, cured mice from C027 treatment rejected subcutaneous flank tumors on rechallenge. These results suggest C027-treatment facilitates establishment of long-term immune memory providing protection from tumor exposure at local and distant sites. Prior studies have shown IL-27 upregulates expression of genes involved in T cell memory differentiation and CD8 T cell intrinsic IL-27 signaling is essential for generating T cell memory responses to subunit vaccination.^46, 47^ While the initial therapeutic effect depends on CTLs, how C027 elicits this observed systemic anti-tumor rejection and if CD8 T cells are participating remains to be clarified. Importantly, the durability and systemic nature of the immune response observed with C027-treatment indicate its potential clinical utility as a therapeutic.

While pre-clinical data is essential for developing new therapeutics, these models have limitations. Syngeneic murine models enable assessment of endogenous immune responses to interventions. However, relative to humans, mice are less susceptible to HSV infection and this can reduce viral replication and gene expression duration.^37^ IL-27 expression and virus replication may differ in human subjects. However, because this cytokine was associated with improved survival in oHSV Phase Ib subjects, this and the murine mechanism identified suggests that it has therapeutic potential. Though IL-27 enhances CTL function and C027-treatment is associated with increased multifunctional CTLs in the glioma TME, the mechanism by which IL-27 mediates these CTL effects requires further investigation. Our results show that these multifunctional CD8 T cells have increased GZB and IFNγ production and suggesting enhancement of a direct cytotoxic function but how IL-27 mediates these effects remains to be answered. Studies have suggested that IL-27 may be altering survival of these cells rather while others found IL-27 may be promoting their proliferation and clonal expansion.^22, 48–50^

In summary, we have developed a novel IL-27-expressing oHSV and demonstrate for the first time oHSV-IL-27 expression enhances survival in syngeneic glioma bearing mice and elicits durable immune memory. Mechanistically, the C027 enhances anti-glioma efficacy in a CD8 T cell and IL-27 dependent manner with associated increases in multifunctional CD8 T cells. Given the promise of current oHSV trials, our studies suggestion local oHSV IL-27 expression may enhance therapeutic efficacy and thus providing a strong rationale for continued investigation and potential clinical translation of IL-27 expressing oHSV for malignant gliomas.

## Declarations

### Patient consent for publication

Not Applicable

### Ethics approval

All animal studies were approved by the Institutional Animal Care and Use Committee (IACUC) at Nationwide Children’s Hospital (IACUC Protocols AR16-00057, AR21-00145).

### Data availability

Data are available upon reasonable request.

### Authors’ contributions

AKM and KAC conceptualized and designed the experiments, and wrote the manuscript. AKM performed the experiments and conducted analyses. US and KAC constructed the C027 virus. JH and IHA assisted with in vivo intracranial surgeries. IHA and DK assisted with in vivo bioluminescent imaging. JH, IHA, DK, YK, and RD assisted with in vivo immunophenotyping experiments. All authors contributed to editing of the manuscript and approved the submitted version. Guarantor author: KAC.

## Acknowledgements

We sincerely acknowledge Carla Waddell and all personnel at the Animal Research Core (Nationwide Children’s Hospital Abigail Wexner Research Institute) for their supervision and care of the mice used in these studies. We also thank the personnel at the Small Animal Imaging Facility and Flow Cytometry Core at Nationwide Children’s Hospital Abigail Wexner Research Institute. Schematic diagrams were created using BioRender.com.

## List of abbreviations

BLI: Bioluminescent imaging
BGS: Bovine growth serum
BFA: Brefeldin A
Breg: Regulatory B cell
CNS: Central nervous system
CM: Conditioned media
CTL: cytotoxic CD8+ T lymphocyte
d: Day
DC: Dendritic cell
DN: dual CD4 and CD8 negative
EFF: Effector
GFP: Enhanced Green Fluorescent Protein
FC: Fold change
FDR: False discovery rate
FBS: fetal bovine serum
ffluc: firefly luciferase
FOL-B: Follicular B cell
GC B: Germinal center B cell
GBM: Glioblastoma
GzB: Granzyme B
h: Hour
HCMV: Human cytomegalovirus
IFNγ: interferon gamma
IL: Interleukin
IC: Intracranial
ip: Intraperitoneal
ITS-B: Isotype switched B cells
LTS: Long-term survivors
MG: Malignant gliomas
MFI: Mean fluorescence intensity
MHC: Major histocompatibility complex
mMDSC: Monocytic myeloid derived suppressor cell
MOI: Multiplicity of infection
MZ-B: Marginal zone B cell
NK: Natural killer
NKT: Natural killer T cell
NS: Not significant
oHSV: oncolytic type I herpes simplex virus
PB: Plasmablast
PC: Plasma cell
PFU: Plaque forming unit
stim: Stimulated
Th1: T helper cell type 1
Th: T helper cell
TCRgd: T cell receptor gamma delta
TNFα: Tumor necrosis factor alpha
Treg: Regulatory T cell
TRM: Tissue-resident memory T cell
TIL: Tumor infiltrating leukocytes
TME: Tumor microenvironment
Tx: Treatment
Tr1: Type I T regulatory cells

