## Supplementary Material for "Oncolytic HSV-IL27 expression improves CD8 T cell function and therapeutic activity in syngeneic glioma models"

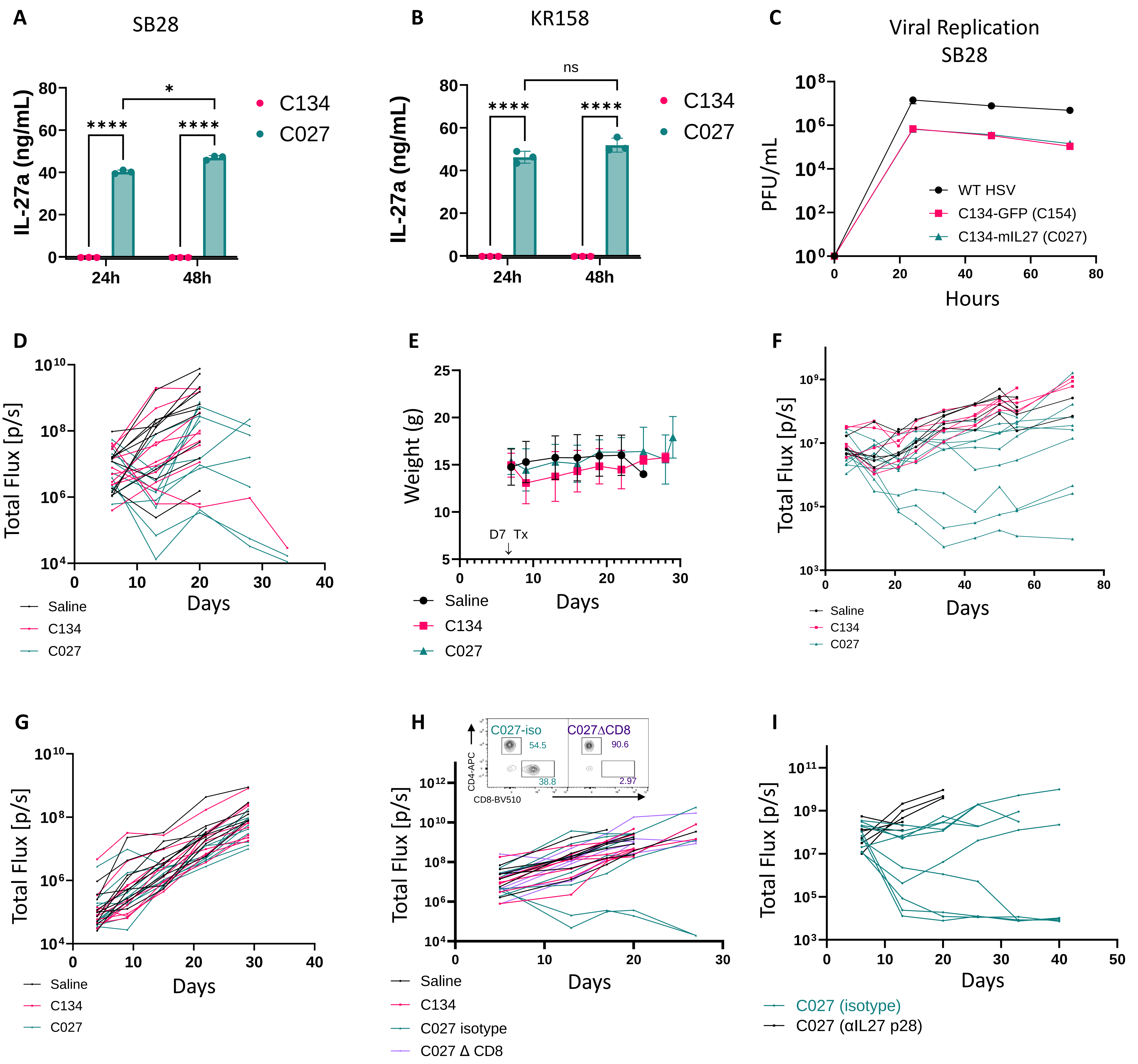


**Supplementary Figure 1**. (A) mIL27 secretion from C027 or C134 infected (MOI=1) in SB28 and (B) KR158 cells. (C) Viral replication kinetics in SB28 cells infected with C027, C134, or WT HSV (MOI=1). Tumor growth curves approximated by bioluminescent imaging (BLI) from mice bearing orthotopic CT-2A (D), KR158 (F), and SB28 (G) gliomas following treatment with saline (black), C134 (pink), or C027 (teal). (E) Mouse weights from mice-bearing orthotopic CT-2A gliomas following treatment. (H) Representative flow plot of peripheral blood confirming depletion efficacy and tumor growth curves (BLI) of orthotopic CT-2A-bearing mice depleted of CD8 T cells by intraperitoneal injection (days 5, 6, 7, 14, 21) of anti-CD8a (purple) or isotype control (teal). (I) Tumor growth curves (BLI) of orthotopic CT-2A-bearing, C027-treated mice receiving intraperitoneal injection (days 7, 9, 11) of IL-27p28 neutralizing antibody (black) or isotype control (teal). (A, B, C) Data are mean ± SEM. (B, C) Each shape representing one replicate. Statistical analyses were performed using two-way analysis of variance with Holm-Šídák's correction for multiple comparisons. Multiplicity of infection, MOI; PFU, plaque forming units.

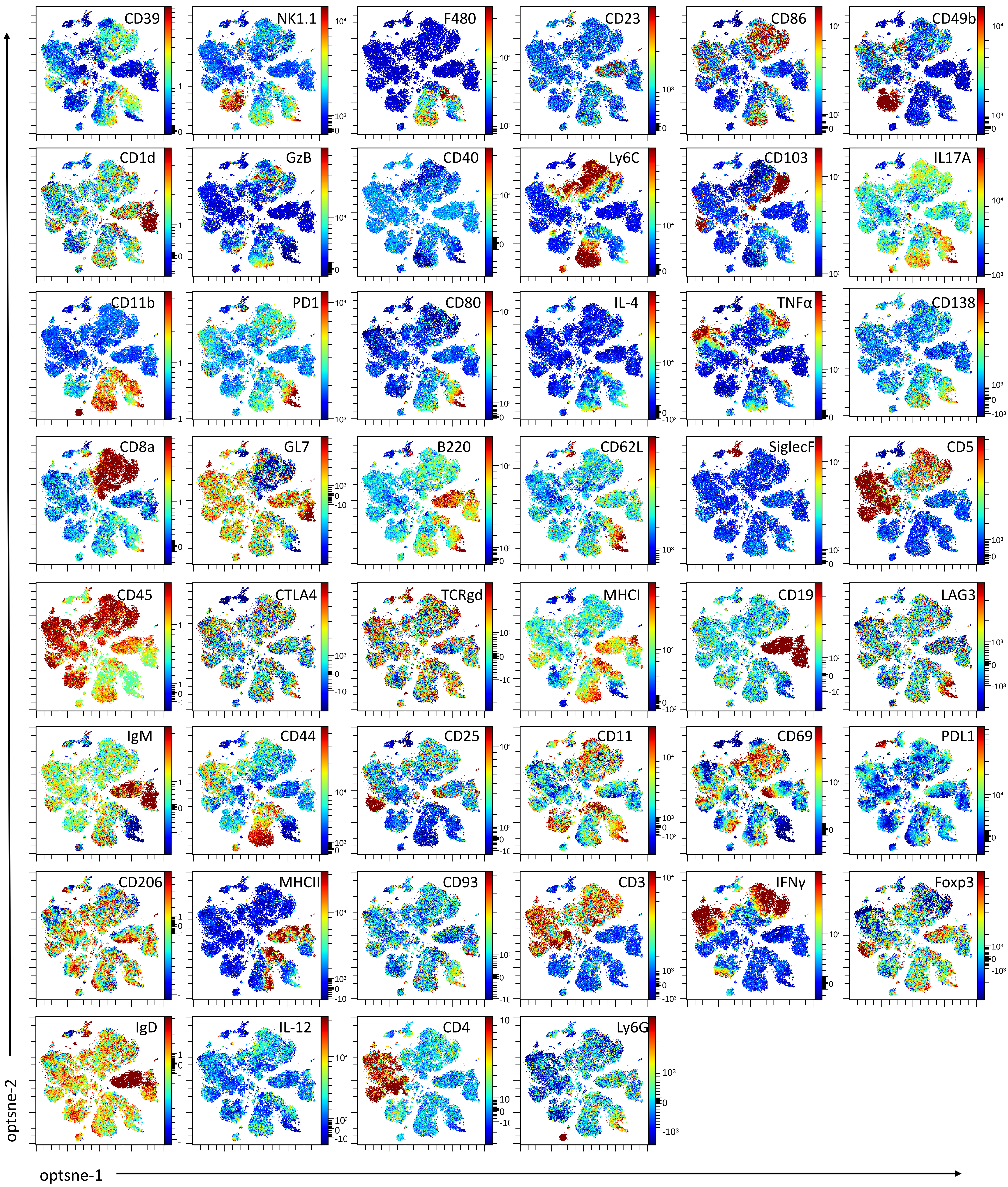


**Supplementary Figure 2**. Spectral Flow Cytometry panel marker expression overlaid on dimensionality reduced opt-SNE plots from CD45+ tumor infiltrating leukocytes. Each point represents a single cell from the concatenated dataset. Color gradient scale represents expression level of each marker. GzB, granzyme B; IFNγ, interferon gamma; TNFα, tumor necrosis factor alpha; MHC, major histocompatibility complex; TCRgd, T cell receptor gamma delta.

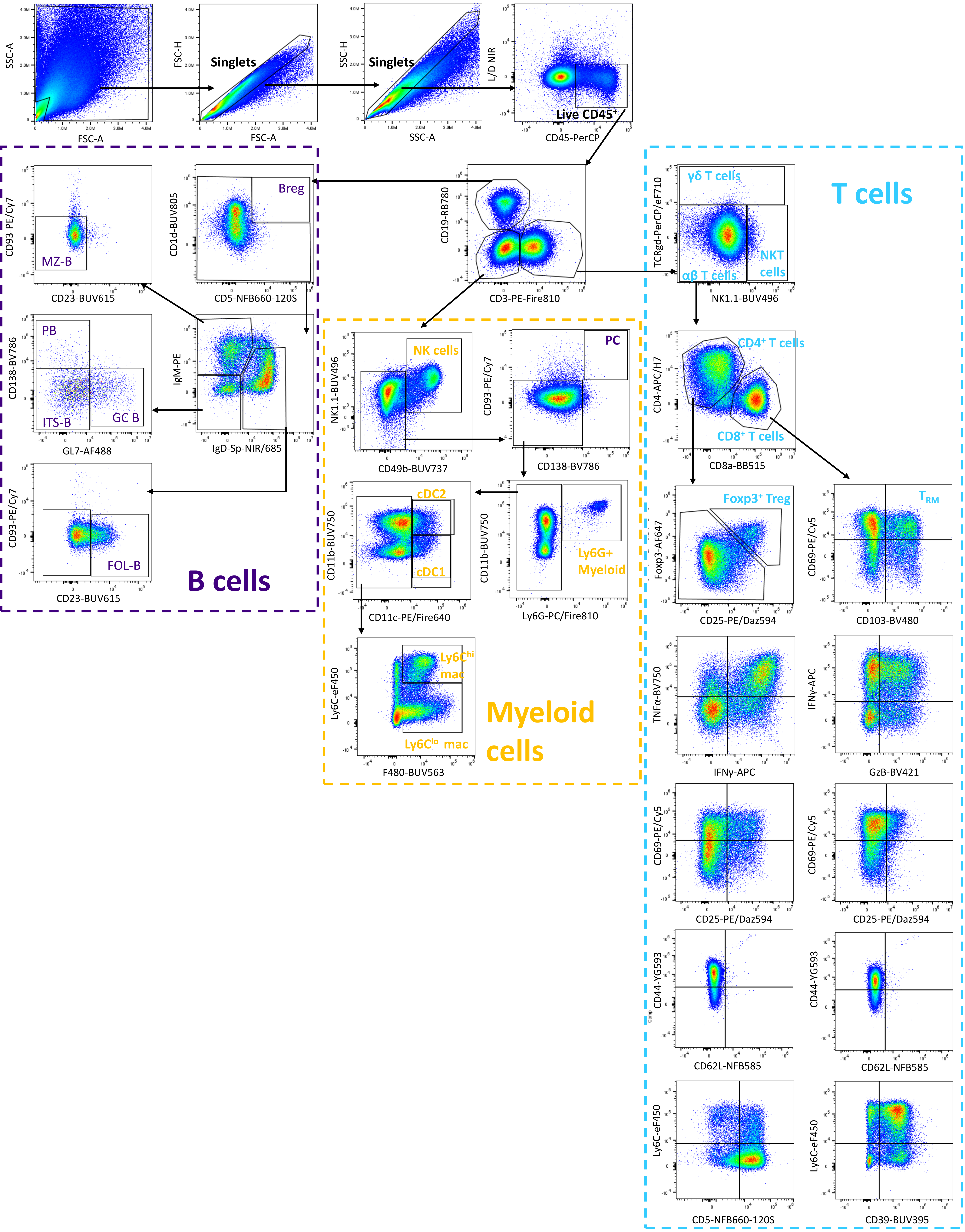


**Supplementary Figure 3**. Flow cytometry gating strategy. Biaxial flow plots depict concatenated live CD45+ cells from tumor infiltrating leukocytes of all samples. Breg, regulatory B cell; PC, plasma cell; PB, plasmablast; ITS-B, isotype switched B cells; MZ-B, marginal zone B cell; FOL-B, follicular B cell; GC B, germinal center B cell; mMDSC, monocytic myeloid derived suppressor cell; DC, dendritic cell; NK, natural killer; NKT, natural killer T cell; TRM, tissue-resident memory T cell; Treg, Regulatory T cell; GzB, granzyme B; IFNγ, interferon gamma; TNFα, tumor necrosis factor alpha.

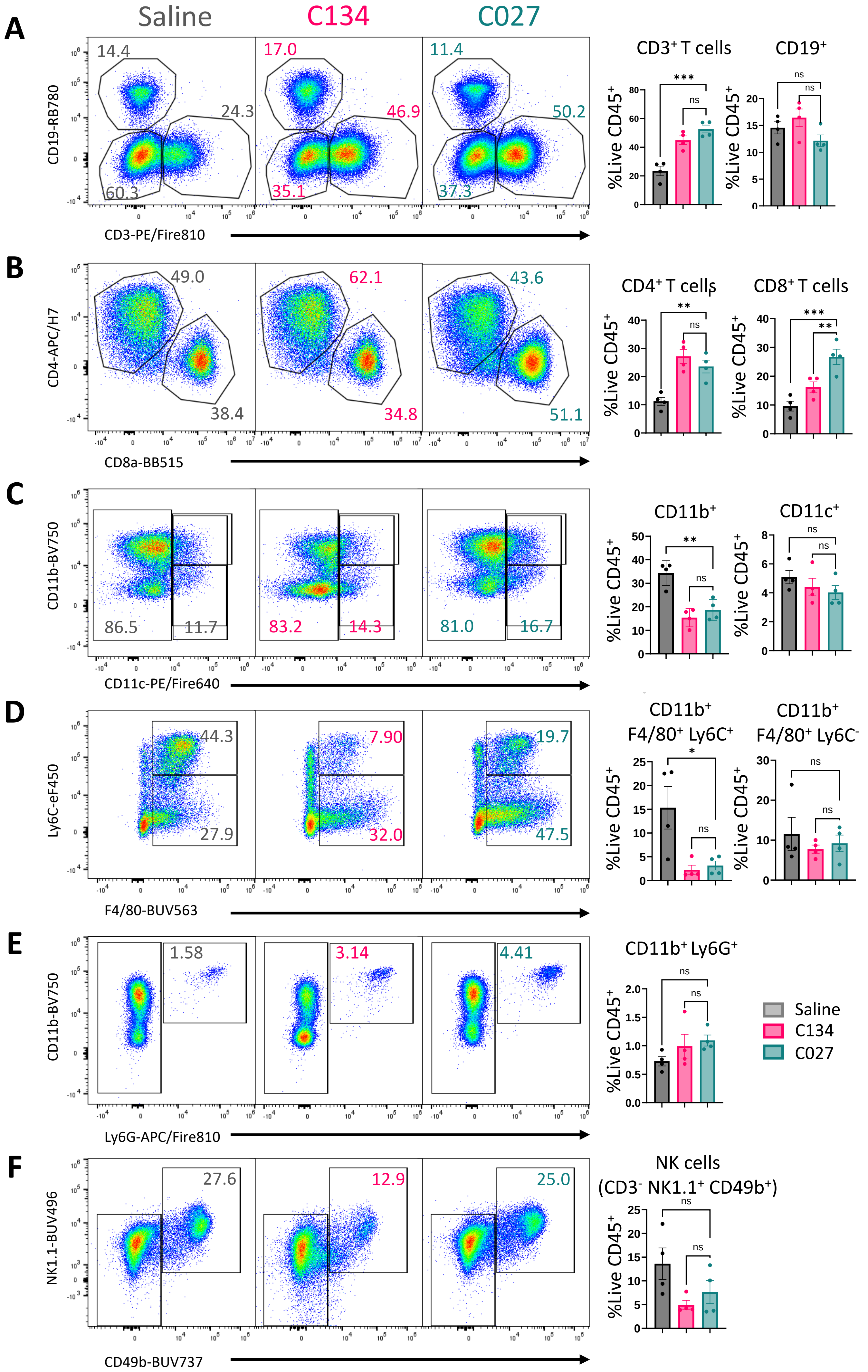


**Supplementary Figure 4**. Flow cytometry gating of immune cell subsets from CD45$^+$ tumor-infiltrating leukocytes. Biaxial flow plots from concatenated live CD45+ cells for each treatment cohort. Frequencies of indicated populations among total CD45$^+$ cells by treatment cohort (n=4 mice per cohort) and depicted as mean ± SEM. Statistical analysis was performed by one-way analysis of variance with Holm-Šídák's multiple comparisons test. NK, natural killer.

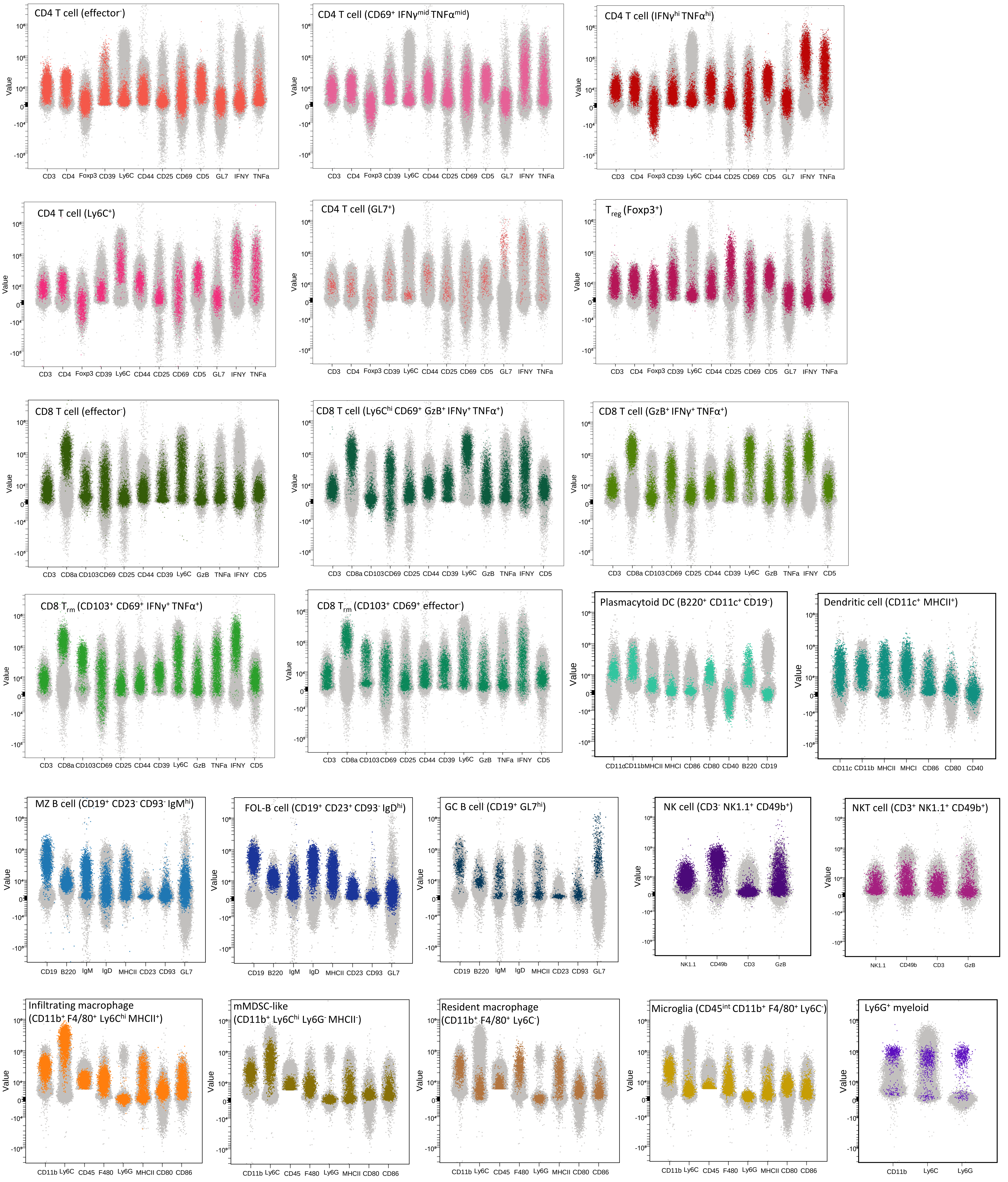


**Supplementary Figure 5**. Scatterplots depicting key marker expression for metaclusters from CD45+ tumor infiltrating leukocytes. Data depicts all concatenated samples for every cluster (gray) or specified cluster (colored). Each point represents one cell and fluorescence value correlates with expression level. Treg, Regulatory T cell; TRM, tissue-resident memory T cell; DC, dendritic cell; MZ-B, marginal zone B cell; FOL-B, follicular B cell; GC B, germinal center B cell; NK, natural killer; NKT, natural killer T cell; mMDSC, monocytic myeloid derived suppressor cell; GzB, granzyme B; IFNγ, interferon gamma; TNFα, tumor necrosis factor alpha; MHC, major histocompatibility complex.

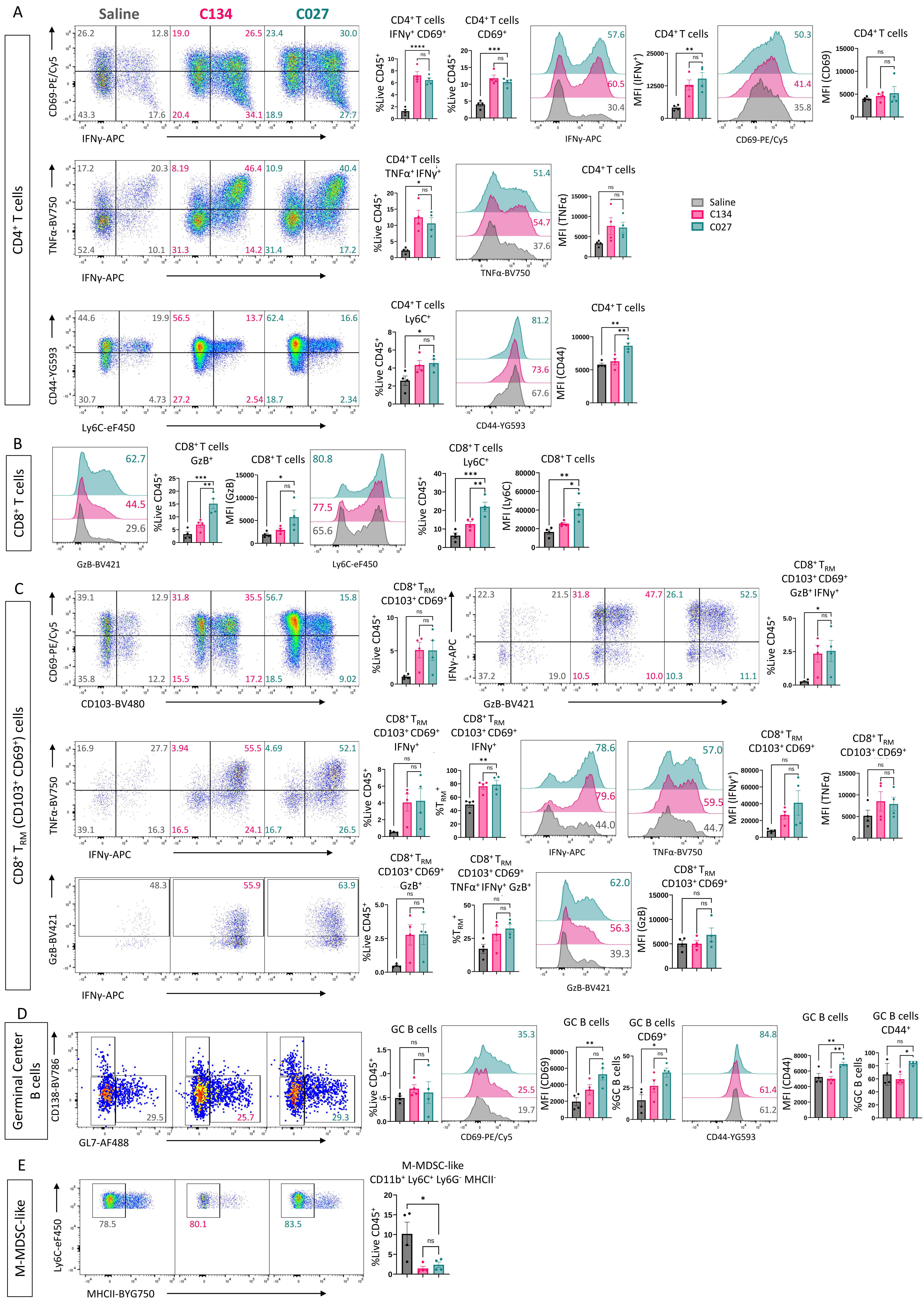


**Supplementary Figure 6**. Flow cytometry gating of T cell subsets from CT-2A tumor infiltrating leukocytes. Biaxial flow plots and histograms depict concatenated live CD45+ cells for each treatment cohort. Frequencies of specified populations among total CD45+ cells by treatment cohort (n=4 mice per cohort) unless otherwise indicated. Expression level (MFI) of indicated markers depicted for specified populations. Data presented as mean ± SEM. Statistical analysis was performed by one-way analysis of variance with Holm-Šídák's correction for multiple comparisons. GC B, germinal center B cell; mMDSC, monocytic myeloid derived suppressor cell; GzB, granzyme B; IFNγ, interferon gamma; TNFα, tumor necrosis factor alpha; TRM, tissue-resident memory T cell; MFI, mean fluorescence intensity; NS, not significant.

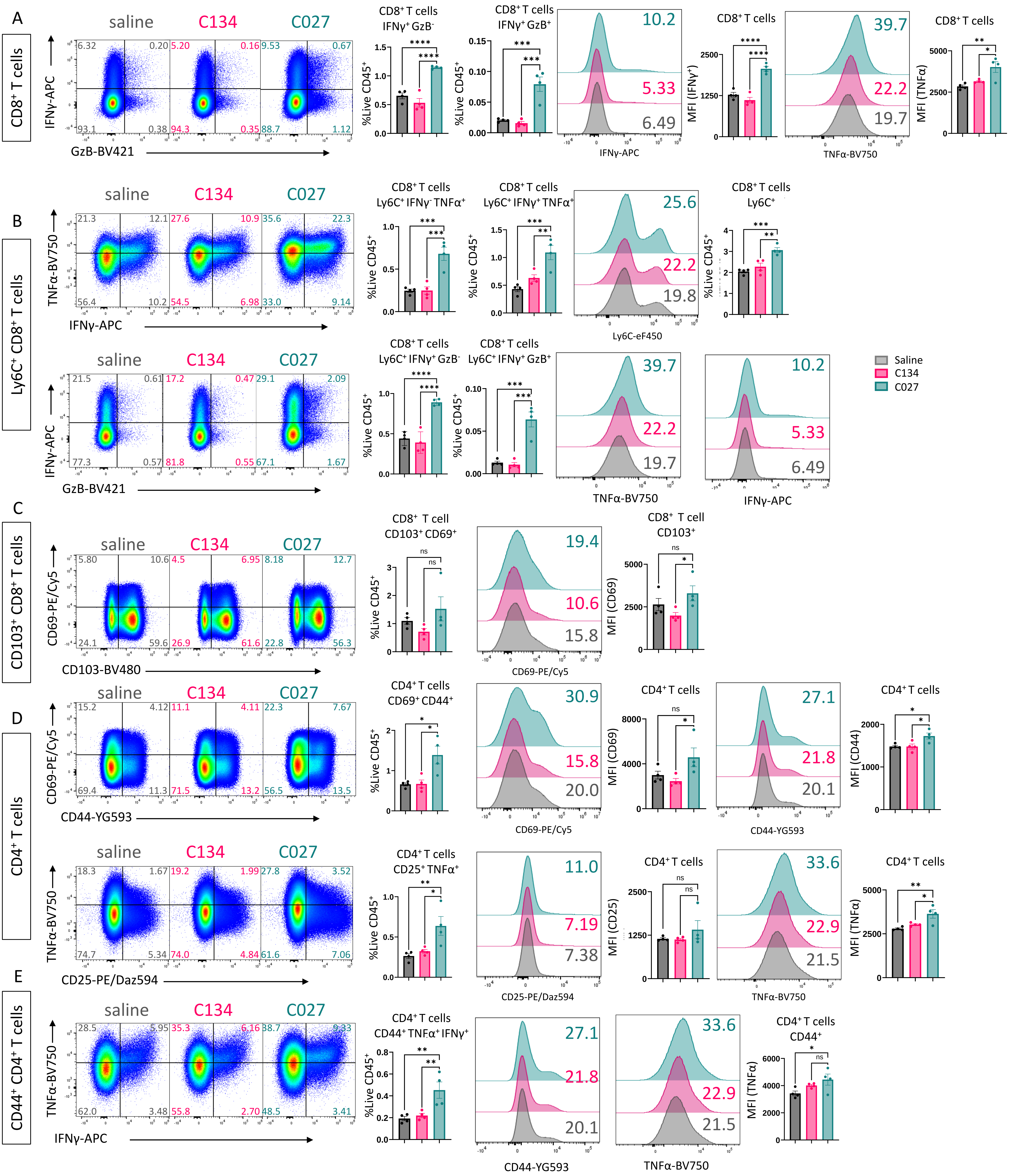

**Supplementary Figure 7.** Flow cytometry gating of T cell subsets from CT-2A spleens. Biaxial flow plots depict concatenated live CD45+ cells for each treatment cohort. Frequencies of specified populations among total CD45⁺ cells by treatment cohort (n=4 mice per cohort) unless otherwise indicated. Statistical analysis was performed by one-way analysis of variance with Holm-Šídák's correction for multiple comparisons. MFI, mean fluorescence intensity; GzB, granzyme B; IFNγ, interferon gamma; TNFα, tumor necrosis factor alpha.

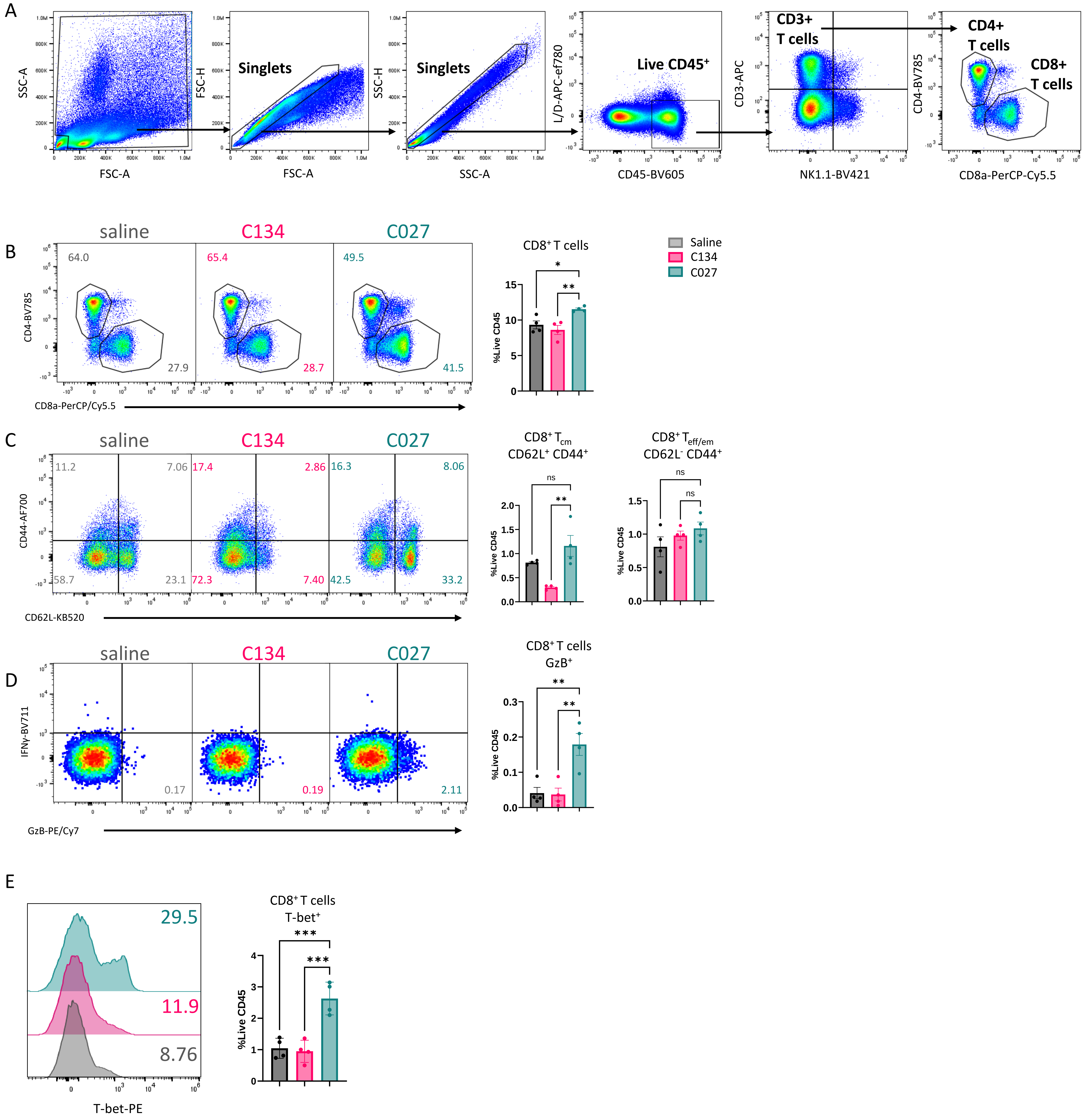


**Supplementary Figure 8**. Flow cytometry gating of T cell subsets from CT-2A-bearing mouse blood Day 12 post-treatment. Biaxial flow plots depict representative flow plots for each treatment cohort. Frequencies of specified populations among total $CD45^+$ cells by treatment cohort (n=4 mice per cohort) unless otherwise indicated. Statistical analysis was performed by one-way analysis of variance with Holm-Šídák's correction for multiple comparisons. Panel: Fixable Viability Stain 780, IFNγ-BV711, GzB-PE/Cy7, T-bet-PE, Foxp3-PE-CF594, NK1.1-BV421, CD45-BV605, CD69-BV650, CD4-BV785, CD62L-KB520, CD8a-PerCP-Cy5.5, CD3-APC, CD44-AF700. Teff/em, Effector/effector memory T cells; Tcm, central memory T cells; T-bet, T-box transcription factor.

**Supplementary Table 1. Spectral flow cytometry antibodies used in these studies.**

| Marker | Conjugate | Clone | Dilution (1 in X) | Company | Catalog # | Staining |
|---|---|---|---|---|---|---|
| Live/Dead | Zombie NIR (Live Dead) | | 800 | Biolegend | 423105 | viability |
| GzB | BV421 | QA18A28 | 400 | Biolegend | 396414 | intracellular |
| IL-17A | eFluor506 | eBio17B7 | 600 | eBioscience | 69-7177-82 | intracellular |
| IL-4 | BV711 | 11B11 | 500 | BD Horizon | 564005 | intracellular |
| TNF-α | BV750 | MP6-XT22 | 400 | Biolegend | 506358 | intracellular |
| IFN-γ | APC | XMG1.2 | 1200 | Biolegend | 505810 | intracellular |
| FoxP3 | AF647 | MF-14 | 500 | Biolegend | 126408 | intracellular |
| IL-12/23 p40 | APC-RF700/AF700 | C15.6 | 500 | Biolegend | 505214 | intracellular |
| CD39 | BUV395 | 723-1185 | 400 | BD Horizon | 567264 | surface |
| NK1.1 | BUV496 | PK136 | 400 | BD OptiBuild | 741062 | surface |
| F4/80 | BUV563 | T45-2342 | 1000 | BD OptiBuild | 749284 | surface |
| CD23 | BUV615 | B3B4 | 2000 | BD OptiBuild | 751169 | surface |
| CD86 | BUV661 | PO3 | 800 | BD OptiBuild | 741528 | surface |
| CD49b | BUV737 | HMα2 | 2000 | BD OptiBuild | 741752 | surface |
| CD1d | BUV805 | 1B1 | 1600 | BD OptiBuild | 741965 | surface |
| CD40 | SB436 | 1C10 | 400 | eBioscience | 62-0401-82 | surface |
| Ly6C | eFluor 450 | HK1.4 | 800 | eBioscience | 48-5932-82 | surface |
| CD103 | BV480 | M290 | 400 | BD Horizon | 566118 | surface |
| CD11b | BV570 | M1/70 | 1200 | Biolegend | 101233 | surface |
| PD1 (CD279) | BV605 | 29F.1A12 | 400 | Biolegend | 135220 | surface |
| CD80 | BV650 | 16-10A1 | 800 | BD Horizon | 563687 | surface |
| CD138 | BV786 | 281-2 | 800 | BD OptiBuild | 740880 | surface |
| CD8a | BB515 | 53-6.7 | 400 | BD Horizon | 564422 | surface |
| GL7 | Alexa Fluor 488 | GL7 | 400 | Biolegend | 144612 | surface |
| B220 (CD45R) | Spark Blue 550 | RA3-6B2 | 800 | Biolegend | 103266 | surface |
| CD62L | NovaFluor Blue 585 | MEL-14 | 400 | eBioscience | M006T02B04 | surface |
| Siglec F (CD170) | Nova Fluor Blue 610-70S | 1RNM44N | 600 | eBioscience | M042T03B06 | surface |
| CD5 | NovaFluor Blue 660-120S | 53-7.3 | 1000 | eBioscience | M026T02B08 | surface |
| CD45 | PerCP | 30-F11 | 2000 | Biolegend | 103130 | surface |
| CTLA4 (CD152) | PerCP/Cy5.5 | UC10-4B9 | 1000 | Biolegend | 106316 | surface |
| TCR γδ | PerCP-eflour 710 | eBioGL3 | 1000 | eBioscience | 46-5711-82 | surface |
| MHC-I (H2-Kb/H2-Db) | RB744 | 28-8-6 | 1000 | BD OptiBuild | 757264 | surface |
| CD19 | RB780 | 1D3 | 1000 | BD OptiBuild | 755522 | surface |
| LAG-3 (CD223) | PerCP/Fire 806 | C9B7W | 500 | Biolegend | 125250 | surface |
| IgM | PE | RMM-1 | 1200 | Biolegend | 406507 | surface |
| CD44 | Spark YG 593 | IM7 | 1000 | Biolegend | 103078 | surface |
| CD25 | PE-Dazzle 594 | PC61 | 400 | Biolegend | 102048 | surface |
| CD11c | PE-Fire 640 | QA18A72 | 600 | Biolegend | 161104 | surface |
| CD69 | PE-Cy5 | H1.2F3 | 600 | Biolegend | 104510 | surface |
| PD-L1 (CD274) | NovaFluor Yellow 690 | MIH5 | 1500 | eBioscience | M036T02Y05 | surface |
| CD206 | PE/Fire700 | C068C2 | 2000 | Biolegend | 141741 | surface |
| MHC-II (IA/IE) | cFluor BYG750 | M5/114.15.2 | 600 | Cytek | R7-20564 | surface |
| CD93 | PE-Cy7 | AA4.1 | 600 | Biolegend | 136506 | surface |
| CD3 | PE-Fire 810 | 17A2 | 800 | Biolegend | 100277 | surface |
| IgD | Spark NIR 685 | 11-26c.2a | 2000 | Biolegend | 405750 | surface |
| CD4 | APC-H7 | GK1.5 | 800 | BD Pharmingen | 560181 | surface |
| Ly-6G | APC-Fire 810 | 1A8 | 1200 | Biolegend | 127670 | surface |

**Supplementary Table 2. Flow cytometry antibodies used in these studies.**

| Marker | Conjugate | Clone | Dilution (1 in X) | Company | Catalog # | Staining |
|---|---|---|---|---|---|---|
| Fixable Viability Stain 780 | APC-ef780 | | 1000 | BD Horizon | 565388 | viability |
| IFN-γ | BV711 | XMG1.2 | 400 | Biolegend | 505835 | intracellular |
| GzB | PE/Cy7 | QA16A02 | 400 | Biolegend | 372213 | intracellular |
| T-bet | PE | 4B10 | 400 | Biolegend | 644809 | intracellular |
| Foxp3 | PE-CF594 | MF23 | 400 | BD Horizon | 562466 | intracellular |
| NK1.1 | BV421 | PK136 | 400 | Biolegend | 108732 | surface |
| PD1 (CD279) | BV510 | 29.F1A12 | 400 | Biolegend | 135241 | surface |
| CD45 | BV605 | 30-F11 | 600 | Biolegend | 103140 | surface |
| CD69 | BV650 | H1.2F3 | 400 | Biolegend | 104541 | surface |
| CD4 | BV785 | GK1.5 | 600 | Biolegend | 100453 | surface |
| CD4 | APC | RM4-5 | 500 | Biolegend | 100516 | surface |
| CD62L | KIRAVIA Blue 520 | MEL-14 | 600 | Biolegend | 104464 | surface |
| CD8a | PE/Cy5.5 | 53-6.7 | 600 | Biolegend | 100734 | surface |
| CD8a | BV510 | 53-6.7 | 500 | Biolegend | 100751 | surface |
| CD25 | AF594 | PC61 | 400 | Biolegend | 102045 | surface |
| CD3 | APC | 17A2 | 600 | Biolegend | 100236 | surface |
| CD3 | FITC | 17A2 | 500 | Biolegend | 100204 | surface |
| CD90.2 | APC | 30-H12 | 600 | Biolegend | 105312 | surface |
| CD44 | AF700 | IM7 | 500 | Biolegend | 103026 | surface |
